# The Multi-allelic Genetic Architecture of a Variance-heterogeneity Locus for Molybdenum Accumulation Acts as a Source of Unexplained Additive Genetic Variance

**DOI:** 10.1101/019323

**Authors:** Simon K.G. Forsberg, Matthew E. Andreatta, Xin-Yuan Huang, John Danku, David E. Salt, Örjan Carlborg

## Abstract

Most biological traits are regulated by both genetic and environmental factors. Individual loci contributing to the phenotypic diversity in a population are generally identified by their contributions to the trait mean. Genome-wide association (GWA) analyses can also detect loci based on variance differences between genotypes and several hypotheses have been proposed regarding the possible genetic mechanisms leading to such signals. Little is, however, known about what causes them and whether this genetic variance-heterogeneity reflects mechanisms of importance in natural populations. Previously, we identified a variance-heterogeneity GWA (vGWA) signal for leaf molybdenum concentrations in *Arabidopsis thaliana*. Here, fine-mapping of this association to a ∼78 kb Linkage Disequilibrium (LD)-block reveals that it emerges from the independent effects of three genetic polymorphisms on the high-variance associated version of this LD-block. By revealing the genetic architecture underlying this vGWA signal, we uncovered the molecular source of a significant amount of hidden additive genetic variation (“missing heritability”). Two of the three polymorphisms on the high-variance LD-block are promoter variants for Molybdate transporter 1 (*MOT1*), and the third a variant located ∼25 kb downstream of this gene. A fourth independent association was also detected ∼600 kb upstream of the LD-block. Testing of T-DNA knockout alleles for genes in the associated regions suggest *AT2G25660* (unknown function) and *AT2G26975* (*Copper Transporter 6; COPT6*) as the strongest candidates for the associations outside *MOT1*. Our results show that multi-allelic genetic architectures within a single LD-block can lead to a variance-heterogeneity between genotypes in natural populations. Further they provide novel insights into the genetic regulation of ion homeostasis in *A. thaliana*, and empirically confirm that variance-heterogeneity based GWA methods are a valuable tool to detect novel associations of biological importance in natural populations.

## Introduction

Genome Wide Association (GWA) analysis is a powerful approach to study the genetic basis of complex traits in natural populations. It is widely used to study the genetics of human disease, but is equally useful in studies of other populations. For example, it has been used to dissect the genetics of traits of importance in agricultural applications (see e.g. [1] for an example in cattle) and ecological adaptation using collections of natural accessions in the genetic model plant *Arabidopsis thaliana* [2–7].

The standard GWA approach screens the genome for loci where the alternative genotypes differ significantly in the mean for the trait or traits of interest. Although hundreds of loci have been found to affect a variety of quantitative traits using this strategy, it has become clear that for most complex traits this additive approach fails to uncover much of the genetics contributing to the phenotypic variation in the populations under study. It is therefore important to explore the genetics of such traits beyond additivity [8]. This could be done by, for example, re-analyzing available datasets to uncover contributions to the phenotypic variability in populations by genetic interactions or variance-heterogeneity [8]. Such analyses have the potential to identify novel loci and alternative genetic mechanisms involved in shaping the diversity in the analyzed populations.

The genetic control of trait variance is a topic that has been studied for many years in quantitative genetics with a primary focus on its potential contributions to adaptation in natural populations and agricultural selection programs. Theoretical and empirical work has increased our understanding of how genetic control of the environmental variance might contribute to phenomena such as fluctuating asymmetry, canalization and genetic robustness [9,10]. Empirical work now also supports the general principle that genetic control of variance heterogeneity is an inherent feature of biological networks and individual genes (see [11] for a review) and that it contributes to both capacitation [12,13] and maintenance of developmental homeostasis [14]. Although it was already shown in the 1980s that it was possible to map Quantitative Trait Loci (QTL) affecting the variance [15], only recently has this approach been more widely adopted to explore the role of variance-heterogeneity loci (or vQTL [16]) in, for example, environmental plasticity [14], canalization [17], developmental stability [18], and natural variation in stochastic noise [19].

With the advent of GWA analysis, and the later realization that standard additive models leave much of the genetic variance in the analyzed populations uncovered [8], there has been an increased interest in exploring genetic regulation of variance heterogeneity [16,20]. Several recent studies in, for example, humans [21,22], plants [6,19,23], *Drosophilia melanogaster* [24] and yeast [25] have shown that part of this previously unexplored heritable genetic variation beyond the narrow-sense heritability can be uncovered by re-analyzing existing GWA datasets using methods to detect differences in trait variance (variance-heterogeneity GWA or vGWA for short) between genotypes [20,22,26]. Despite these findings that variance-controlling loci appear to be relatively common in the genome and can make large contributions to the phenotypic variability [23,25], the molecular basis of variance-controlling loci remain to be uncovered.

Further, much work remains to empirically evaluate the validity of the wide variety of hypotheses that have been proposed regarding the origin of the variance-heterogeneity signals. These include variance-heterogeneity being a single locus property due to for example allelic heterogeneity, incomplete linkage disequilibrium and developmental instabilities [6,16,23], or the result of its interactions with other genetic or environmental factors (i.e. epistasis or gene-by-environment interactions) [16,21,25]. The first suggestive empirical evidence for variance-heterogeneity being a property of a single locus was recently presented in an unpublished study by Ayroles *et al.* [24]. In that study, a vGWA association to a behavioral phenotype in the Dropsophila Genome Reference Panel (DGRP) was determined to be driven by a mutation in the *Ten-a* gene. However, more work is needed to fully understand the molecular genetic mechanisms underlying variance-heterogeneity associations to evaluate how these loci contribute to the genetic architecture of complex traits.

Allelic heterogeneity could cause a genetic variance-heterogeneity in a GWA study where tests for associations are to bi-allelic markers (see e.g. [16]). It is well known that major loci affecting traits under selection often evolve multiple mutations with variable effects on the phenotype. Some examples of this have been studied in detail in domestic animal populations, and major contributions have been illustrated for Mendelian traits, such as coat color [27–29], and complex traits including muscularity in cattle [30] and meat quality in pigs [31]. Allelic diversity has also been found in natural populations, where the work by Poormohammad *et al.* [32] is particularly interesting for this study as it illustrates the potentially adaptive value of allelic diversity at the Molybdate transporter 1 (*MOT1*) gene in *A. thaliana*. Despite this, complex trait genetic studies often ignore such allelic complexity in the initial genome-wide analyses although some exceptions exist [33,34].

The vGWA approach opens new opportunities to identify such multi-allelic loci via a straight-forward extension of the standard GWA framework [20]. The reason for this can be illustrated using the following example. Consider a GWA analysis to a bi-allelic SNP marker that tags two versions of an LD-block in the genome. If within this LD-block more than two functional alleles exist that contribute to the trait of interest, the SNP marker will not be able to tag these perfectly. If the functional alleles are evenly distributed across the two versions of the LD block tagged by the SNP genotypes, there will be no association detected in the GWA regardless of whether the test is performed on the scale of the mean or the variance. If, on the other hand, the functional alleles are unevenly distributed such that the version of the LD-blocked tagged by one of the SNP marker alleles is associated with multiple functional variants with different effects on the trait, whereas the version tagged by the other marker allele is associated with functional alleles with the same phenotypic effect, there will be a difference in variance between the genotypes. If all of the alleles associated with one of the genotypes either increase or decrease the phenotype when compared to the allele at the other locus, some effect can also be observed on the mean. If, on the other hand, some of the alleles increase and others decrease the trait relative to the allele on the other version of the LD-block, only a variance effect can be observed. Hence, re-analyses of GWA datasets using a vGWA approach, followed by explorations of the underlying mechanisms, present new opportunities for identifying contributions by allelic heterogeneity to the variability of complex traits in natural populations.

Previously, we re-analyzed ionomic data from an early trait-mean based GWA study that had used 93 wild-collected *A. thaliana* accessions [2] and detected a variance-heterogeneity GWA with genome-wide significance for variation in leaf molybdenum concentrations. This association is located near the *MOT1* (Molybdate transporter 1) gene [23]. Importantly, this locus did not affect the mean leaf molybdenum concentrations in either of these studies [2,23]. Molybdenum is an essential element for plant growth due to its role as a part of the molybdopterin cofactor that is required by several critical enzymes [35]. Both deficiency and excess of Molybdenum have an impact on plant development [36]. The ability of plants to acquire minerals from the soil, and regulate their levels in the plant, requires complex biochemical and regulatory pathways. The genetic architecture of such ionomics traits is thus complex [37]. To date, only a limited number of studies in *A. thaliana* have exploited natural variation and QTL analysis to examine mineral content [38–44], and only nine that have identified the molecular determinants of the QTL. These include Co, Mo, Na, Cd, As, S/Se, Zn, Cu and sulfate [5,7,32,45–52]. GWA have also been used to identify both candidate loci and functional polymorphisms contributing to natural variation in these traits [2,3,5,7,53].

Here, we use data for molybdenum concentrations in leaves of more than 300 natural *A. thaliana* accessions for which genome-wide genotypes were publicly available [3]. Using this data, we replicate the genetic variance heterogeneity association previously detected by Shen *et al.* [23] in a different set of accessions. Further, we show that this genome-wide association results from a complex multi-locus, multi-allelic genetic architecture in a ∼78 kb segment surrounding the *MOT1* locus. Variation in leaf molybdenum concentrations is associated with three loci in this segment. In two of these, the minor alleles are structural polymorphisms in the *MOT1* promoter region. The third is a SNP in LD with a functional candidate gene (*AT2G25660*) identified to affect leaf molybdenum concentrations through analysis of a T-DNA insertion allele. A fourth associated SNP was located ∼600 kb from *MOT1* and in LD with the *Copper Transporter 6* (*AT2G26975*/ *COPT6*) gene for which an evaluated T-DNA insertion allele had decreased leaf molybdenum concentrations. *COPT6* is an interesting candidate gene as copper and molybdenum homeostasis has recently been shown to be connected [54]. This connection may in part be due to the need for copper during the biosynthesis of molybdopterin [55].

We demonstrate how multi-allelic genetic architectures can lead to vGWA associations and illustrate the type of multi-allelic complexity that can evolve at loci determining adaptively important phenotypes in natural populations. Further, dissection of variance-heterogeneity loci reveals novel additive genetic variance that otherwise would remain undetected and contribute to the “missing heritability”.

## Results

### An increased population-size reveals novel GWA associations affecting Molybdenum concentrations in leaves

The first GWA analysis searching for genetic effects on the mean leaf molybdenum concentrations [2] failed to uncover any genome-wide significant associations for this trait. This was surprising as it was known from earlier QTL studies that a strong polymorphism affecting this trait was segregating in this population [46]. To investigate this further we measured the leaf molybdenum concentration in leaves from at least six replicate plants of 340 natural *A. thaliana* accessions (S1 Table) that had earlier been genotyped using the 250k *A. thaliana* SNP-chip [3]. In this larger and more powerful dataset, several GWAs to the mean leaf molybdenum concentration are revealed to be in, or near, the *MOT1* locus (Fig 1). The minor alleles for some associated SNPs increased the mean phenotype, whereas others decreased it relative to the major allele (Table 1; Fig 1B; p_raw, top SNP_ = 3.1 × 10^-14^). Importantly, in an earlier study, we identified a genome-wide significant genetic variance-heterogeneity association between leaf molybdenum concentrations and a locus containing *MOT1* [23]. We therefore used statistical and functional approaches to dissect this region further to obtain a deeper understanding of the genetic mechanisms controlling these different types of natural variation in leaf molybdenum we observe in *A. thaliana*.

**Table 1.** Mean and variance effects for five loci in the *MOT1* region associated with either mean molybdenum concentration levels (GWA) or variance (vGWA).

| Locus | SNP<sup>1</sup> | Location (bp)<sup>2</sup> | Effect (95% CI)<sup>3</sup> | Significance<sup>4</sup> |
| --- | --- | --- | --- | --- |
| **Mean associations** |  |  |  |  |
| DEL<sub>1</sub> | n.a. | 10,934,564 - 10,934,616 | -2.56 (-3.16 - -1.96) | $4.2 \times 10^{-16}$ |
| DUP<sub>1</sub> | n.a. | 10,934,814 - 10,935,143 | 2.75 (1.95 - 3.55) | $5.0 \times 10^{-11}$ |
| SNP<sub>1</sub> | rs347469902 | 10909091 | 0.82 (0.42 - 1.22) | $4.6 \times 10^{-5}$ |
| SNP<sub>2</sub> | rs347287517 | 11528777 | 0.99 (0.69 - 1.29) | $3.0 \times 10^{-10}$ |
| **Variance association** |  |  |  |  |
| vBLOCK<sub>1</sub> | rs346812256 | 10928720 | 2.69 (2.20 - 3.29) | $4.4 \times 10^{-10}$ |

<sup>1</sup>Name of the most strongly associated SNP markers at each locus in the GWA (Mean associations) or vGWA (Variance association) analyses; <sup>2</sup>Location in the TAIR10 assembly ; <sup>3</sup>Estimates of mean effects ( $\mu\text{g Mo /g dry weight}$ ) and 95% Confidence Intervals (CI) from a four-locus fixed-effect regression model including DEL<sub>1</sub>, DUP<sub>1</sub>, SNP<sub>1</sub> and SNP<sub>2</sub>, and the variance effect (fold increase in variance for the high-variance genotype) for vBLOCK<sub>1</sub> from a dGLM model;
<sup>4</sup>Significances for the effects estimated in<sup>3</sup>

**Fig 1.**
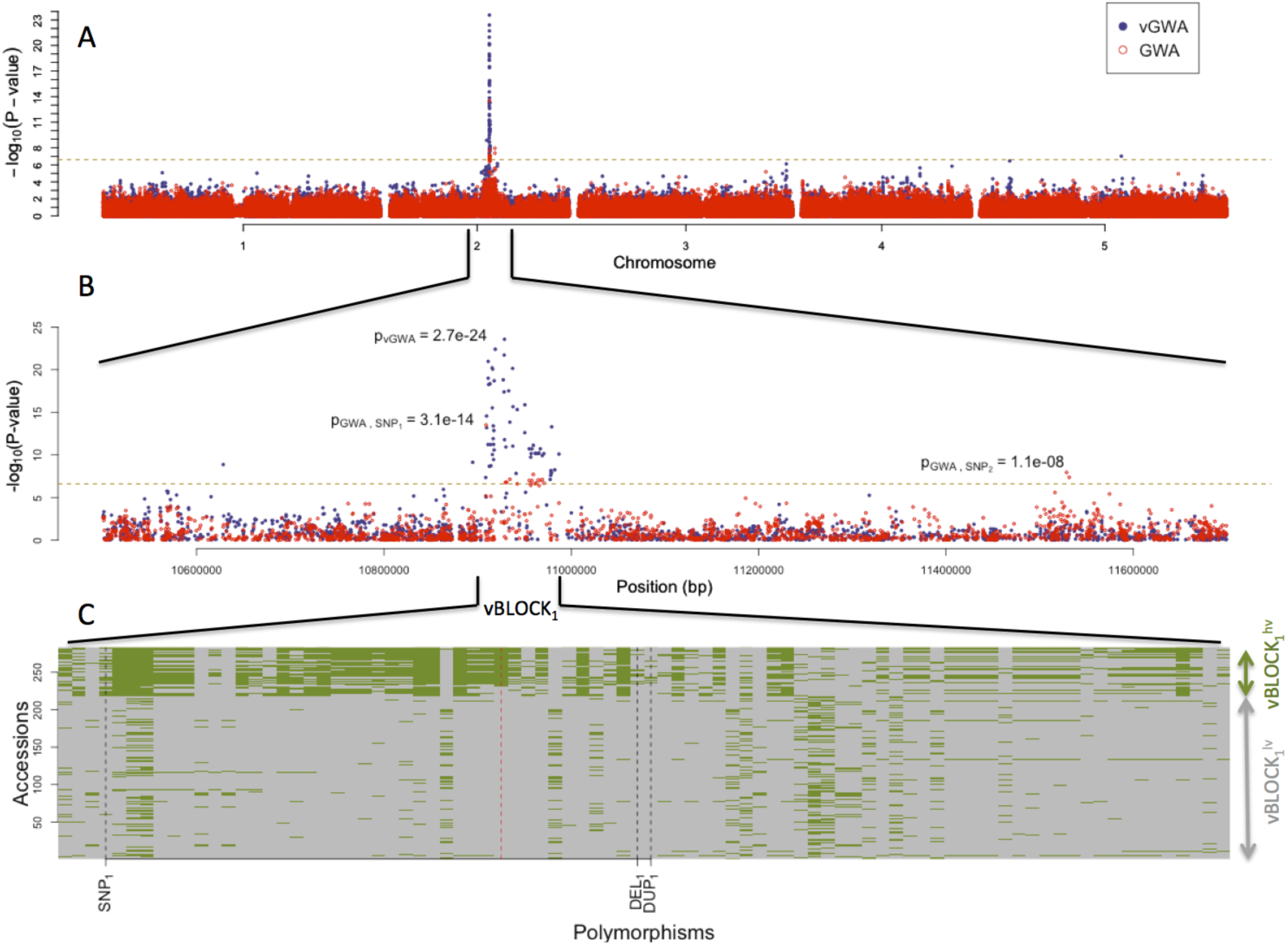
GWA and vGWA analyses for mean leaf molybdenum concentration. (A) Genome-wide results from single-locus vGWA (blue) and GWA (red) analyses across the *A. thaliana* genome. (B) Region on chromosome 2 with the strongest vGWA association to the mean leaf molybdenum concentrations. The LD is high between a large number of SNPs across a ∼78 kb region (vBLOCK_1_), where the minor alleles define a high-variance associated version of this chromosomal segment *ν BLOCK*_*1*_^*hv*^. (C) Illustration of the high LD across vBLOCK_1._ The accessions that are homozygous for the minor/major allele are colored green/grey and then sorted according to the genotype of SNP with the strongest association in the vGWA (Table 1).

### *Dissecting the genetic structure of a variance-heterogeneity locus affecting molybdenum concentrations in* A. thaliana

A high-resolution vGWA analysis of leaf molybdenum concentrations in the 340 accessions revealed several genome-wide significant associations that overlapped with the earlier reported signal near the *MOT1* gene [23]. These associations were very strong (Fig 1A; p_raw, top vSNP_ = 1.55 × 10^-27^) at a number of SNPs in a ∼78 kb LD-block on chromosome 2 (Fig 1B; 10, 908,677-10,986,893 bp; vBLOCK_1_). By visualizing the multilocus genotypes for the analyzed accessions across vBLOCK_1_, we observed that the population contains two distinct multi-locus genotype classes for this segment: one that predominantly contains high-variance associated SNP alleles *ν BLOCK*_*1*_^*hv*^ and another with low-variance associated SNP alleles (*ν BLOCK*_*1*_^*lv*^; Fig 1C). vBLOCK_1_ contains in total 20 annotated genes, and the most obvious functional candidate for the association is *MOT1* (10,933,061-10,934,551).

One possible explanation for the vGWA signal we observe is what is known as allelic plasticity. Here, alternative alleles at a locus respond differently to, for example, environmental or genetic perturbations. Allelic plasticity can be separated from other potential mechanisms underlying genetic variance heterogeneity as it is the only one leading to both within- and between-genotype variance-heterogeneity, whereas the other only result in a between-genotype variance-heterogeneity [16]. As our dataset contained multiple replicates for each accession, we could test for the presence of a within-genotype variance heterogeneity, quantified as the coefficient of variation (CV) of molybdenum concentrations, across the vBLOCK_1_ region (see Methods section for further details). No genetic effect on the CV was found in our data, and based on this we conclude that allelic plasticity is an unlikely explanation for the vGWA we observe in this region.

### *A multi-locus genetic architecture contributes to the variability in molybdenum concentrations in* A. thaliana

Multi-allelic genetic architectures can lead to a genetic variance-heterogeneity [16]. To illustrate this, let us use the following simple theoretical example in an inbred (or haploid) organism. First assume that the locus under study contain three functional alleles (A, B and C) with A being the major allele, and B and C minor alleles that increase and decrease the phenotype, respectively, relative to the major allele. A standard GWA analysis associates the phenotype with the genotypes at bi-allelic SNP markers. When the true genetic architecture is multi-allelic, no perfect associations will exist between the SNP marker genotypes and the functional alleles. Such imperfect associations will often lead to a variance-heterogeneity between the SNP genotype classes that can be detected via a vGWA analysis. Consider, for example, when alleles B and C are in LD with each other and one of the SNP marker alleles, whereas allele A is in LD with the other SNP allele. Then there is no mean difference between the genotypes, but all the genetic variance is instead captured as a variance-heterogeneity between the genotypes (S1 Fig). The phenotypic distribution for the high-variance SNP allele becomes larger as it is a mixture between those of alleles B and C (S1 Fig). The following sections show that the vGWA to vBLOCK_1_ is largely due to such a multi-allelic genetic architecture.

#### Multiple structural MOT1 promotor-polymorphisms are associated with molybdenum concentrations in leaves

*MOT1* is an obvious functional candidate gene for the vGWA to vBLOCK_1_. A 53 bp deletion in the promoter-region of the gene has earlier been shown to decrease *MOT1* expression, leading to low concentrations of molybdenum in the plant [46,56]. To complement our SNP-marker dataset with this known, and other potentially functional, structural promoter polymorphisms segregating in the analyzed population, we screened the promoter region of *MOT1* and identified five loci and six non-coding structural polymorphisms (Fig 2, S2 Table) that were then genotyped in 283 of the 340 phenotyped accessions.

**Fig 2.**
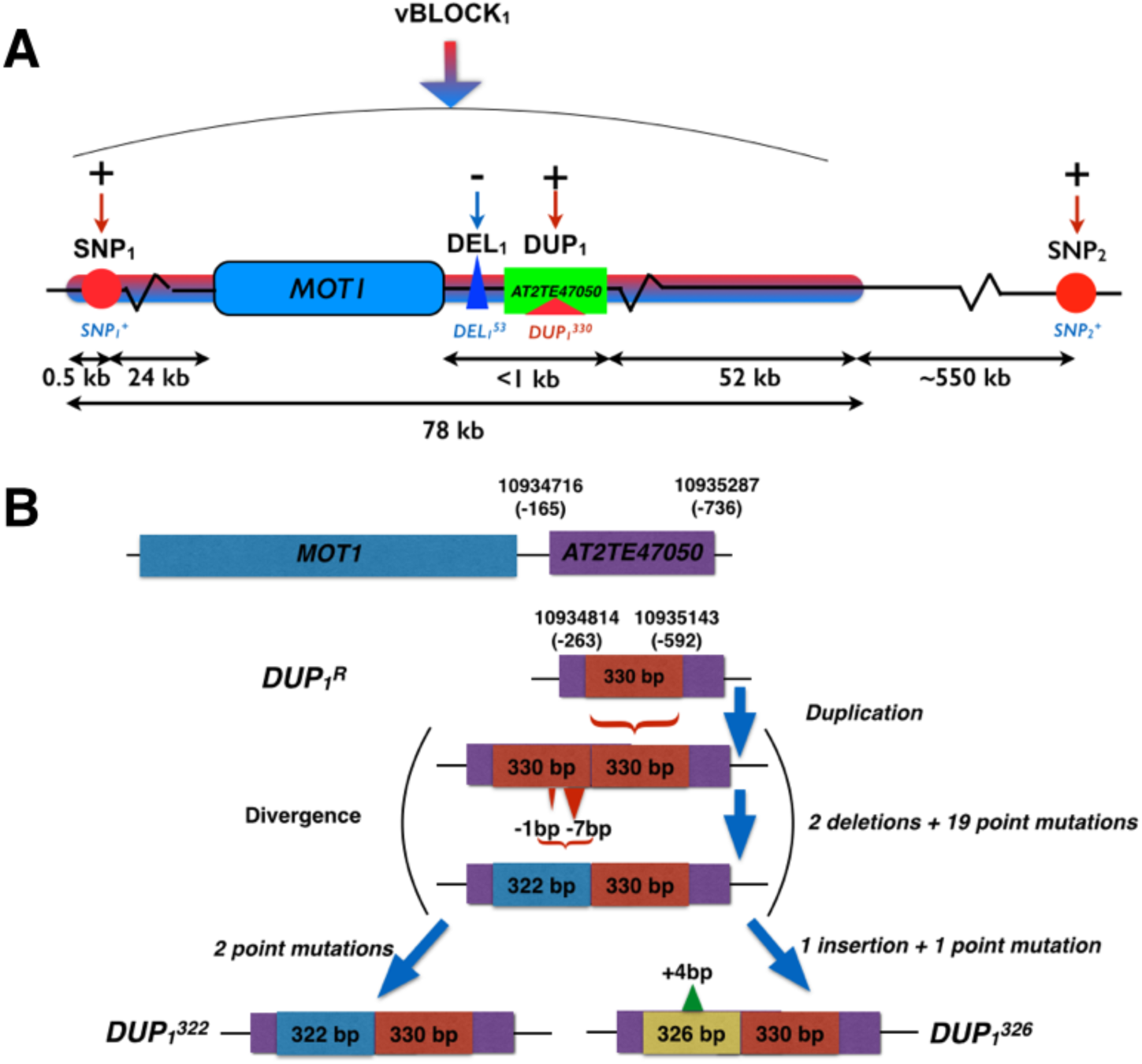
Schematic illustration of GWA and vGWA signals for mean leaf molybdenum concentrations. (A) A vGWA signal was detected to an ∼78 kb region surrounding *MOT1* (Red/blue arrow; vBLOCK_1_). Three GWA signals were detected within (SNP_1_, DEL_1_ and DUP_1_) and a fourth (SNP_2_) upstream of vBLOCK_1._ The direction of the effects for the minor alleles at these loci (*SNP*_*1*_^*+*^, *SNP*_*2*_^*+*^, *DEL*_*1*_^*53*^ and *DUP*_*1*_^*326*^) relative to that of the major, reference allele are illustrated with + (increased) and - (decreased), respectively. In (B) we illustrate the differences between the reference allele at DUP_1_ (*DUP*_*1*_^*R*^) and the two variants of the 330 bp duplication (*DUP*_*1*_^*326*^ and *DUP*_*1*_^*322*^) in the transposable element *AT2TE47050* in the promoter region of *MOT1*.

Two of the six segregating *MOT1* promoter polymorphisms altered the mean leaf molybdenum concentration. The first was *DEL*_*1*_^*53*^ which is located 13 bp upstream from the transcription start-site of *MOT1*. Baxter *et al.* [46] earlier showed that this 53 bp deletion (*DEL*_*1*_^*53*^) allele lacks the TATA-box in the *MOT1* promoter which leads to a reduced expression of *MOT1* and decreased molybdenum concentration in the leaf. We confirm that this allele decreased the mean molybdenum concentrations in the leaf also in this dataset (Table 1; p_raw_ = 9.3x10^-11^; Fig 2A) and found the *DEL*_*1*_^*53*^ allele only among low molybdenum accessions (Mo < 3ppm). An even stronger association p_raw_ = 1.4x10^-15^; Table 1; Fig 2A) was found to a locus (DUP_1_) located 263 bp upstream from the translation start site. Here, several accessions share a 330bp long duplication (Fig 2B) located inside a transposable element (*AT2TE47050*). The duplication exists in two distinct variants (alleles) differing by four polymorphisms: three point-mutations and one 4bp insertion (*DUP*_*1*_^*326*^ and *DUP*_*1*_^*322*^ in Fig 2B). In our dataset, the *DUP*_*1*_^*326*^ allele altered leaf molybdenum concentrations and it was found only among accessions with high leaf molybdenum concentrations (Mo > 10ppm). These observations provide further evidence that allelic heterogeneity at *MOT1* is an important component of the genetic architecture of natural variation in leaf molybdenum concentrations.

#### A multi-locus analysis confirms that a multi-locus, multi-allelic genetic architecture determines the variation in molybdenum concentrations in plants from the global A. thaliana *population*

Multiple associations to the mean- and variance of mean leaf molybdenum concentrations were uncovered in the single-locus analyses. To confirm the independence of these effects, and evaluate their joint contributions to the variation in molybdenum in the population, we fitted all markers (SNPs and structural variants) on chromosome 2 in a generalized linear model to the mean leaf molybdenum concentration using the Lasso method [57]. This penalized maximum likelihood regresses the effects of polymorphisms that make no, or only a minor, independent contribution to the trait towards zero and highlights the markers that jointly make the largest contribution to the trait variation. The penalty in the analyses was chosen so that all highlighted polymorphisms in the final model also have a genome-wide significant effect in the earlier GWA or vGWA analyses (see Methods section for details). In this way, the Lasso method picks up the genome-wide significant polymorphisms that have independent effects.

The loci DEL_1_ and DUP_1_, where the minor alleles were the structural *MOT1* promoter polymorphisms, *DEL_1_^53^* and *DUP_1_^326^*, were the most strongly associated loci in, or near, the *MOT1* gene in the Lasso analysis. Two additional SNP markers, one located ∼25 kb downstream (rs347469902; 10,909,091 bp; SNP_1_; Table 1) and one ∼600 kb upstream of *MOT1* (rs347287517; 11,528,777 bp; SNP_2_; Table 1), were also highlighted. The minor alleles at SNP_1_ and SNP_2_ (*SNP1+* and *SNP2+*) were both enriched among accessions with high leaf molybdenum concentrations. The minor alleles at three of the four associated loci thus increased the mean leaf molybdenum concentrations (Table 1; *DUP*_*1*_^*326*^, *SNP*_*1*_^*+*^, and *SNP*_*2*_^*+*^),and one decreased it (Table 1; *DEL_1_^53^*).

#### Three polymorphisms affecting mean molybdenum concentrations explain the major variance-heterogeneity association peak

Our vGWA and GWA analyses identify five genome-wide significant associations. One vGWA where many SNPs distributed across an ∼78 kb LD-block were associated with the variance of leaf molybdenum concentrations (vBLOCK_1_). Then four GWA where the minor alleles altered the mean leaf molybdenum concentrations relative to the population mean. The pairwise LD between the four loci affecting the molybdenum mean was low (r^2^ < 0.2; Table 2), as expected given their independent contributions to the trait. In all 20 accessions that carry either the *DEL_1_^53^* or the *DUP_1_^326^* alleles, and in 19 of the 29 accessions carrying the high molybdenum *SNP1+* allele, these alleles were present on the same, high-variance associated multi-locus *ν BLOCK*_*1*_^*hv*^ (Fig 1C; see Methods section for further detail). This results in a high LD (D‘ = 1) between these polymorphisms (*DUP*_*1*_^*326*^, *DEL*_*1*_^*53*^ and *SNP*_*1*_^*+*^) and the high-variance associated alleles across vBLOCK_1_ (Fig 1C; Table 2). The high-variance associated SNPs in vBLOCK_1_ are thus in high LD with multiple alleles that either increase or decrease the mean molybdenum concentration in the plant, suggesting a multi-allelic, multi-locus genetic architecture as an explanation for the variance GWA associations we observe in this region. It is, however, worth noting that the estimates of LD using r^2^ is low (r2=0-0.19), illustrating that this measure is not as useful for identifying polymorphisms that potentially contribute to a multi-allelic vGWAs.

**Table 2.** LD between the loci altering mean leaf molybdenum concentrations. LD is provided as r^2^ /D’ above/below the diagonal, respectively.

|  | **DEL<sub>1</sub>** | **DUP<sub>1</sub><sup>a</sup>** | **SNP<sub>1</sub>** | **SNP<sub>2</sub>** | **vBLOCK<sub>1</sub><sup>b</sup>** |
| --- | --- | --- | --- | --- | --- |
| **DEL<sub>1</sub>** | 1 | 0 | 0.01 | 0 | 0.16 |
| **DUP<sub>1</sub><sup>a</sup>** | 0.96 | 1 | 0.19 | 0.03 | 0.10 |
| **SNP<sub>1</sub>** | 0.99 | 0.99 | 1 | 0.03 | 0.43 |
| **SNP<sub>2</sub>** | 0.59 | 0.52 | 0.22 | 1 | 0.03 |
| **vBLOCK<sub>1</sub><sup>b</sup>** | 0.99 | 0.99 | 0.92 | 0.18 | 1 |

<sup>a</sup>Minor allele in the analysis is *DUP<sub>I</sub><sup>326</sup>* that is associated with mean leaf molybdenum concentrations; <sup>b</sup>vBLOCK1 represented by the top-SNP in the vGWA analysis (Table 1).

To statistically disentangle the genetic effects on the mean and variance by this multi-allelic, multi-locus genetic architecture, an additional vGWA analysis was performed where we fitted a linear model with separate effects for the mean and variance to the data as outlined by Valdar and Rönnegård [16]. The three GWA associated loci that were located within vBLOCK_1_ (DUP_1_, DEL_1_ and SNP_1_) were fitted as loci with mean effects when screening chromosome 2 for loci with potential effects on the variance using this method. The entire vGWA signal to vBLOCK_1_ disappears in this analysis (Fig 3A) illustrating that the variance-heterogeneity association to vBLOCK_1_ is due to the presence of the *DEL_1_^53^*, *DUP_1_^326^* and *SNP_1_^+^* alleles on the high-variance associated *ν BLOCK*_*1*_^*hv*^ (Fig 3C).

**Fig 3.**
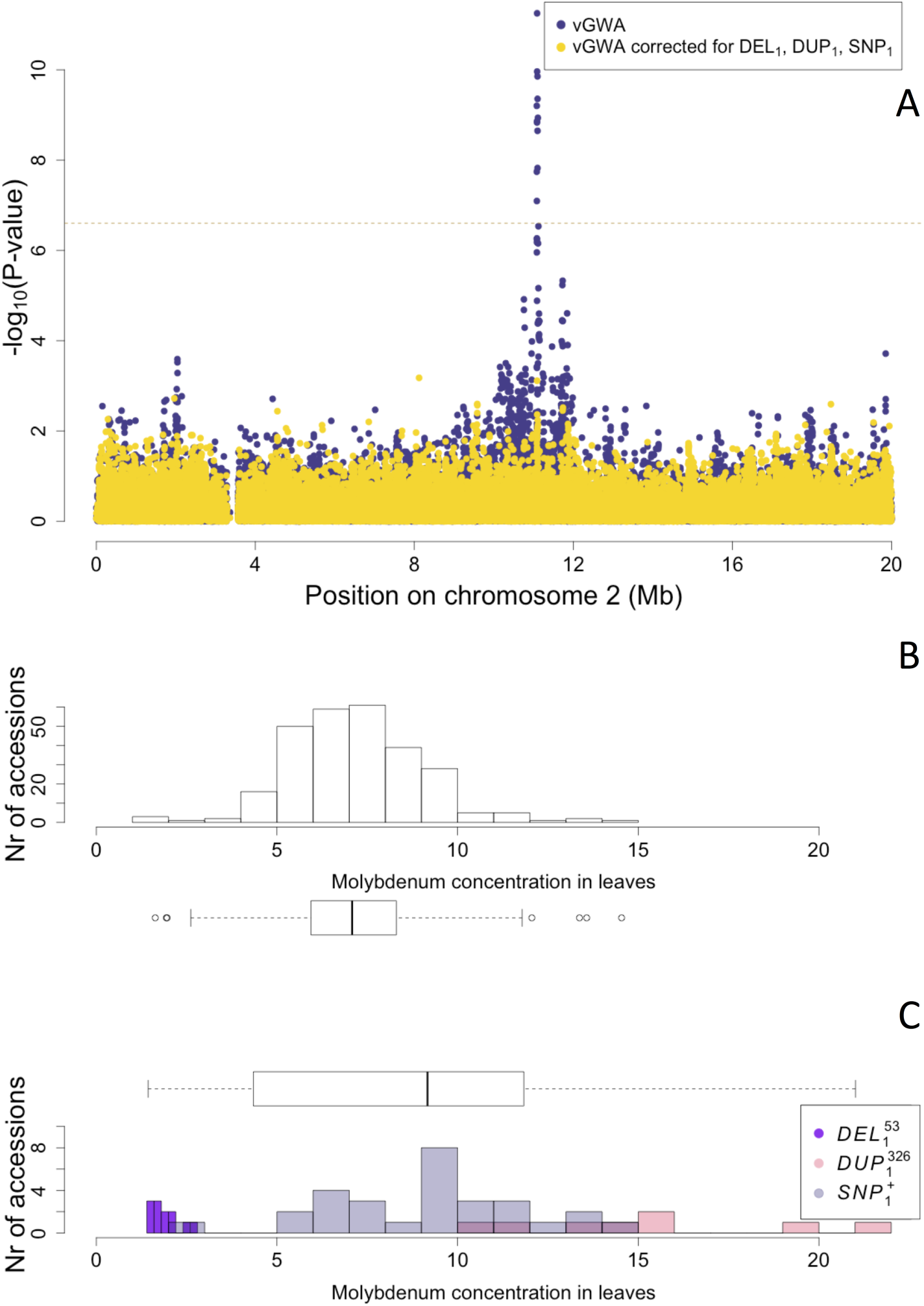
The vGWA signal to vBLOCK_1_ emerges from a multi-locus, multi-allelic genetic architecture. **(A)** The vGWA analysis detects a strong association near *MOT1* on chromosome 2 also when the DGLM approach is used (blue dots). This vGWA signal is, however, cancelled when the mean effects of the *DEL*_*1*_^*53*^, *DUP*_*1*_^*326*^ and *SNP*_*1*_^*+*^ alleles are included in the DGLM model (yellow dots). The variation in mean leaf molybdenum concentrations is lower among the accessions carrying the low-variance associated variant of vBLOCK_1_ *ν BLOCK*_*1*_^*lv*^ **(B)** than among the accessions carrying the high-variance associated variant *ν BLOCK*_*1*_^*hv*^ **(C)**. Separate colors are used for the accessions carrying the *DEL*_*1*_^*53*^ (purple), *DUP*_*1*_^*326*^ (red) and *SNP*_*1*_^*+*^ (grey) alleles in **(C)** to illustrate how these alleles generate the high variance in mean leaf molybdenum concentrations associated with *ν BLOCK*_*1*_^*hv*^.

### New additive genetic variation revealed by the dissection of a vGWA signal

We estimate the broad-sense heritability of leaf molybdenum concentrations from the within/between accession variances to be H^2^= 0.80 using an ANOVA across all replicated measurements, which is similar to that reported in earlier studies (0.56 [53] −0.89 [2]). The narrow-sense heritability was estimated to be h^2^ >=0.63 using a mixed model based analysis where the accession mean phenotypes were regressed onto the genomic kinship matrix.

The first GWA analysis for leaf molybdenum concentrations by Atwell *et al.* [2] was unable to detect any loci contributing to the genetic variation for this trait. The later vGWA study by Shen *et al.* [23] identified a vGWA signal in the *MOT1* region that explained 27% of σP^2^, where the contribution by mean (additive) and variance (non-additive) effects were 4/23% of σP^2^, respectively. Using the variance decomposition proposed by Shen *et al.* [23], we estimate that the vGWA signal to vBLOCK_1_ contributes 3 and 19 % to σP^2^ via its effect on the mean and the variance heterogeneity. The total amount of genetic variance associated with the vGWA signal here is thus comparable to that of Shen *et al.* [23], but in both studies it leaves much of the total additive genetic variance unexplained as it only accounts for about 5% of h^2^. The contribution to H^2^ is, however, larger and between 24 to 28% in these two studies.

However, after considering the individual contributions made by the three polymorphisms identified on *vBLOCK*_*1*_^*hv*^ (*DEL_1_^53^*, *DUP_1_^326^*, *SNP_1_^+^*; Fig 3), much additive genetic variance is uncovered. Nearly all the contribution from vBLOCK_1_ becomes additive (83% of the total variance) to explain 45% of h^2^ and 43% of H^2^. By also accounting for the fourth locus (SNP_2_; Fig 2), the contribution h^2^ and H^2^ increases further to 60 and 50 %, respectively. By dissecting the genetic architecture of the vGWA signal into the contribution by this allelic heterogeneity, we were thus able to reveal a significant contribution by vBLOCK_1_ to the “missing heritability” of molybdenum concentration in the leaf in the original GWA [2] and vGWA [23] analyses.

### Functional analyses of genes in LD with the GWA associations

Here, we functionally explore the associations outside of the coding and regulatory regions of *MOT1* in more detail to identify additional functional candidate polymorphisms and genes for the regulation of molybdenum homeostasis.

#### Mutational analyses to identify functional candidates contributing to variable leaf molybdenum concentrations in A. thaliana

Two regions outside of the coding and regulatory region of *MOT1* (chromosome 2 10,933,061-10,935,200 bp) were associated with the mean leaf molybdenum concentrations (SNP_1_ and SNP_2_ in Fig 1B; 3A). Genes located in the chromosomal regions covered by SNPs in LD (r^2^ > 0.4) with SNP_1_ and SNP_2_, respectively, were explored as potential functional candidates for the associations using T-DNA insertion alleles (S3 Table).

Five T-DNA alleles of five different genes in the region around SNP_1_ (10,909,091 bp; Fig 4A; S3 Table) were evaluated for leaf molybdenum concentrations. One line (SALK_048770) with an insertion in *EMBRYO DEFECTIVE 2410* (*AT2G25660;* 10,916,203 - 10,927,390 bp) displayed a significant 65% decrease in leaf molybdenum concentrations compared to the wild-type (p = 0.018; Fig 4; Table 3). The function of this gene is largely unknown, but is required for normal embryo development [58] and is differentially expressed in pollen [59]. No relationship to molybdenum has been reported previously.

**Table 3.** T-DNA insertion lines with significant associations to the mean leaf molybdenum Concentrations.

| T-DNA Line | ATG Number | Gene | Molybdenum concentration<br>($\mu\text{g} / \text{g}$ dry weight)<sup>a</sup> | p-value<sup>b</sup> |
| --- | --- | --- | --- | --- |
| SALK_048770 | AT2G25660 | EMBRYO DEFECTIVE 2410 | 10.6 (30.4) | 0.018 |
| SALK_138758 | AT2G27020;AT2G27030 | PAG1;CAM5 | 3.5 (36.3) | 0.014 |
| GK-350E02 | AT2G26975 | COPT6 | 0.6 (4.4) | 0.007 |

<sup>a</sup>Mean leaf molybdenum concentration ( $\mu\text{g} / \text{g}$ dry weight; n=6) for the tested T-DNA insertion line (*Col-0* reference) on the tray where it was tested ; <sup>b</sup>Significance as determined by Wilcox rank test comparing molybdenum concentration in the leaves of the T-DNA insertion line to that of the *Col-0* reference on the same tray.

**Fig 4.**
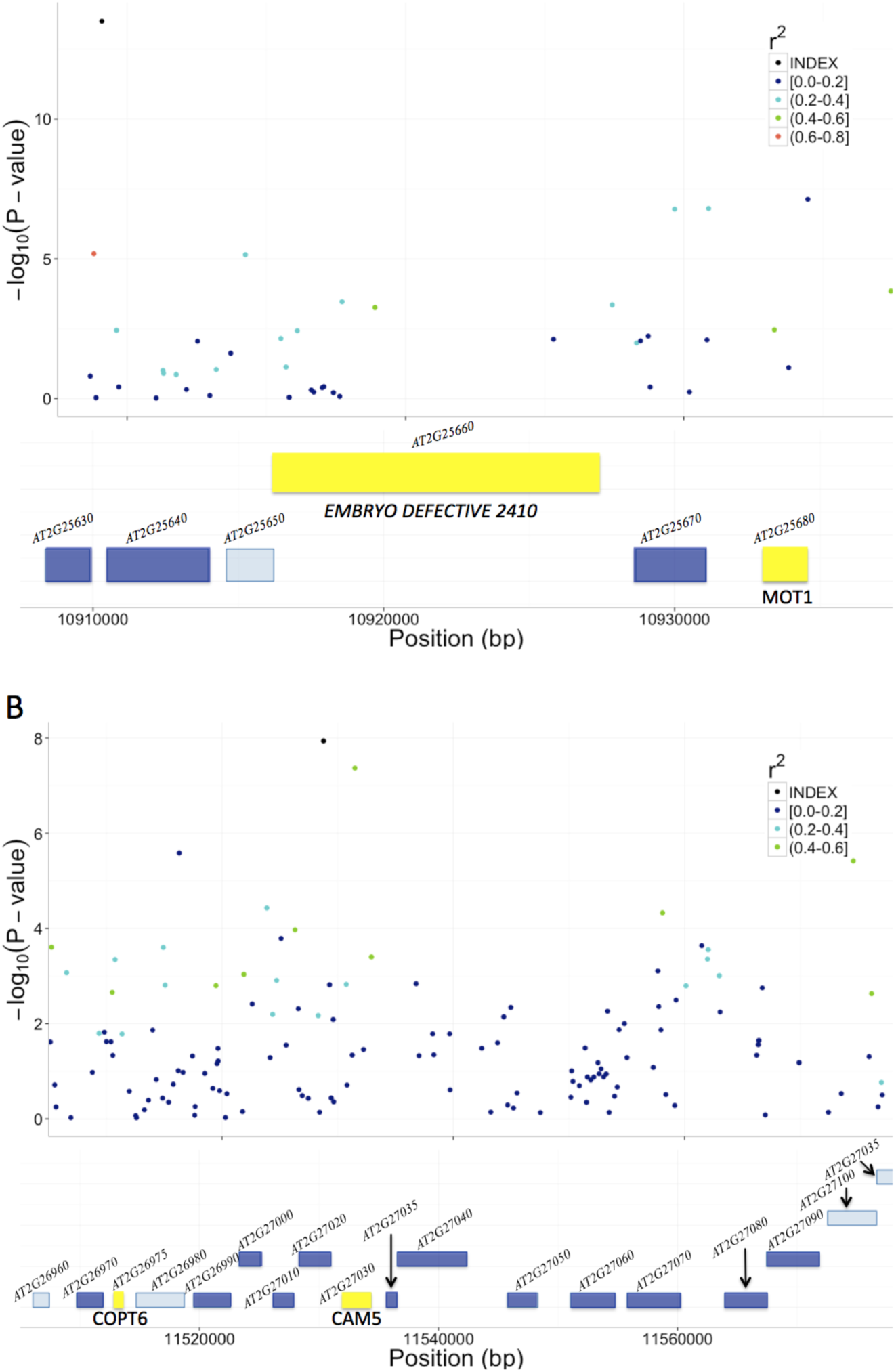
T-DNA analyses to identify candidate genes for the associations to mean leaf molybdenum concentrations. We identified the genes (colored boxes) in the regions surrounding the SNP_1_ and SNP_2_ GWA that were bounded by the furthest up- and downstream SNPs with r^2^ > 0.4. We measured the mean leaf molybdenum concentrations for available T-DNA insertion lines and compared them to the reference genotype (*Col-0*). Yellow box = significant difference in leaf molybdenum concentration, deep blue box = no significant difference, light blue = no T-DNA insertion line tested. **(A)** In the region surrounding SNP_1_, a single T-DNA line with an insertion in *AT2G25660* had an altered mean leaf molybdenum concentration. **(B)** In the region surrounding SNP_2_, the T-DNA lines with insertions in *AT2G26975* and between *AT2G27020/AT2G27030* had altered mean leaf molybdenum concentrations.

We also evaluated 19 T-DNA lines with insertions in 14 genes around SNP_2_ (11,528,777 bp; Fig 4B; S3 Table), and identified two with significantly altered leaf molybdenum concentrations compared to the wild-type (Table 3). One (SALK_138758) was an insertion covering *AT2G27020* and *AT2G27030*, and the other (GK-350E02) had an insertion in *AT2G26975*. These T-DNA alleles showed 90 and 86% reductions in leaf molybdenum concentrations compared to wild-type, respectively (Table 3). *AT2G27020* was also evaluated via another T-DNA insertional allele (SAIL_760_D06), and this line had wild-type leaf molybdenum concentrations. Thus, *AT2G27030* (*ACAM2*/*CAM5*; 11,532,004 - 11,534,333) appears to be the most likely functional candidate gene of the two. Calmodulin is a known metalloprotein and a Ca^2+^ sensor, but no previous connections to molybdenum has been reported. Based on its reduced leaf molybdenum *AT2G26975* (Copper Transporter 6; *COPT6*) is a second functional candidate locus for the association around *SNP*_*2*_. Interestingly, as well as low molybdenum, the T-DNA knockout allele of this gene has a slightly increased leaf copper concentration compared to wild-type (3.82 and 3.36 µg / g dry weight, respectively, in GK-350E02 and *Col-0*; p = 0.0018), suggesting a role of *COPT6* also in the regulation of copper homeostasis. From the literature it is known that copper and molybdenum homeostasis are related and that copper depleted *Brassica napus* plants have upregulated expression of both copper transporter genes and *MOT1* [54].

## Discussion

Common approaches to dissect the genetics of complex traits in segregating populations are linkage mapping and association studies. These studies aim to identify the loci in the genome where genetic polymorphisms regulate the phenotypic variability in the studied populations. This is achieved by screening for significant genotype-phenotype associations across a large number of genotyped polymorphic markers in the genome. The most common statistical models used in such analyses aim to identify loci with significant mean phenotype differences between the genotypes at individual loci, i.e. additive genetic effects. Although additive models are powerful for capturing much genetic variance in populations, they have limited power when challenged with more complex genetic architectures including multiple-alleles, variance-heterogeneity and genetic interactions [8,60]. It is therefore important to also develop, and test, methods that explore statistical genetic models reaching beyond additivity when aiming for a more complete dissection of the genetic architecture of complex traits.

The genetic architecture of mean leaf molybdenum concentrations has earlier been explored using GWA analyses in a smaller set of 93 wild collected *A. thaliana* accessions [2]. No genome-wide significant associations were found, which was surprising given that the trait has a high heritability [46,53] and that several *MOT1* polymorphisms are known to contribute to natural variation in this trait [32,46]. We later found that a non-additive genetic mechanism - genetic variance-heterogeneity - explained a large fraction of the phenotypic variance for a range of phenotypes in the Atwell *et al.* data [23] and the strongest association for leaf molybdenum concentrations was found near the *MOT1* gene. Here, we dissect the genetic architecture underlying this strong association using a larger set of 340 accessions. Due to the higher power of this dataset, we were able to split this high-variance association into four novel GWAs to the mean concentration of leaf molybdenum. The minor allele at one of these (*DEL_1_^53^*) was a deletion in the promoter region of *MOT1* previously identified using an F_2_ bi-parental mapping population. This deletion allele decreases the concentration of molybdenum in leaves by down-regulating *MOT1* transcription [46]. Three novel associations were also found and the minor alleles at these loci (*DUP_1_^326^*, *SNP_1_^+^* and *SNP_2_^+^*) increased the concentration of molybdenum in leaves. One (*DUP_1_^326^*) was an insertion polymorphism in the promoter region of *MOT1*, and the other two associations to SNPs in regions that were not in LD with the *MOT1* gene or its promoter. One of these SNPs was found ∼25 kb downstream of *MOT1* (SNP_1_) and the other ∼600 kb upstream of the *MOT1* transcription start-site (SNP_2_). The regulation of molybdenum concentrations in the leaves is hence due to multiple alleles in a gene known to regulate molybdenum uptake, *MOT1*, but also alleles at other neighboring loci that have earlier not been found to contribute to molybdenum homeostasis in *A. thaliana*. Our results thus provide the first successful replication of a variance-heterogeneity locus on a genome-wide significance level in an independent dataset. Further, they reveal strong associations in both the vGWA and GWA analyses with distinct overlap with findings from earlier QTL and functional analyses of the *MOT1* region [32,46], stressing the central importance of this region in the regulation of molybdenum homeostasis in natural populations.

Little is known about the genetic mechanisms contributing to variance-heterogeneity between genotypes in natural populations. Recently, Ayroles *et al.* [24] showed in an unpublished study that allelic plasticity at an individual locus could be a cause, but beyond that other explanations are still hypothetical. Here, we find that the variance-heterogeneity in leaf molybdenum concentration revealed in a collection of wild-collected *A. thaliana* accessions results from extensive LD across an ∼78 kb chromosomal region surrounding *MOT1* (vBLOCK_1_). The high-variance associated variant of this block (*ν BLOCK*_*1*_^*hv*^) contains three independent polymorphisms (*DEL_1_^53^*, *DUP_1_^326^* and *SNP_1_^+^*) altering the molybdenum concentration in leaves relative to the major alleles at these loci on the low-variance associated variant (*ν BLOCK*_*1*_^*lv*^).Two of these polymorphisms increase molybdenum and one decrease it, leading to a highly significant genetically determined variance-heterogeneity amongst the accessions that share *ν BLOCK*_*1*_^*hv*^. Our study thus present the first empirical evidence illustrating how population-wide variance heterogeneity in a natural population can result from allelic heterogeneity, and an illustration of how the use of non-additive genetic models in GWA analyses can provide novel insights to the genetic architecture of complex traits in natural populations.

Many GWA studies have found that the total additive genetic variance of associated loci is considerably less than that predicted based on estimates of the narrow-sense heritability, i.e. the ratio between the additive genetic and phenotypic variance in the population. This common discrepancy between the two is often called the curse of the “missing heritability” and is viewed as a major problem in past and current GWA studies [61]. Here, we provide an empirical example of how a vGWA is able to identify a locus [23] that remained undetected in a standard GWA [2] and that, when the underlying genetic architecture was revealed, was found to make a large contribution to the additive genetic variance and narrow-sense heritability. This illustrates the importance of utilizing multiple statistical modeling approaches in GWA studies to detect the loci contributing to the phenotypic variability of the trait, and then also continue to further dissect the underlying genetic architecture to uncover how the loci potentially contribute to the heritability that was “missing” in the original study [2].

Here, we dissected a vGWA association to the molybdenum concentration in *A. thaliana* leaves into an underlying multi-locus, multi-allelic genetic architecture. We find several alleles at *MOT1* that contribute to this association, which is consistent with findings in earlier studies that have found that several functional variants of this gene alter the mean molybdenum concentrations in *A. thaliana* [32,46]. Our results also support the earlier notion that potentially important contributions from vGWA [23] and multi-allelic genetic architectures [33,34]contribute to “missing heritability”. The vGWA is thus an alternative, straight-forward and computationally tractable analytical strategy to identify loci where multi-allelic genetic architectures reduce the additive genetic variance that can be detected by traditional GWA approaches.

By evaluating T-DNA mutant alleles in genes in LD with the SNPs associated to leaf molybdenum concentrations, we are able to suggest three novel functional candidate genes involved in molybdenum homeostasis in *A. thaliana*. Little is known about the function of two of these, *AT2G25660* and *AT2G27030*, and further work is needed to explore the mechanisms by which they may alter molybdenum concentrations in the plant. The fourth gene (*AT2G26975*; Copper Transporter 6; *COPT6*) located ∼600 kb upstream of *MOT1* is from earlier studies known to be involved in the connected regulation of copper and molybdenum homeostasis in plants. It was recently reported [54] that *MOT1* and several copper transporters were upregulated under copper deficiency in *B. napus*, suggesting a common regulatory mechanism for these groups of genes. Further experimental work is needed to explore the potential contributions of these genes to natural variation in molybdenum homeostasis and the potential connection between copper and molybdenum homeostais.

In summary, we dissect a vGWA to the leaf molybdenum concentration in *A. thaliana* [23] into the contributions from three independent alleles that are co-localized on the high-variance associated variant of a long (∼78 kb) LD block surrounding the *MOT1* gene. Quantitative genetic analyses show that vGWAs mark additive genetic variation that is undetectable using standard GWA approaches. The dissection of the genetic architecture underlying the vGWA signal allowed the transformation of non-additive genetic variance into additive genetic variance, and hence allowed the detection of a significant part of the “missing heritability” in the variation in leaf molybdenum concentrations in this species-wide collection of *A. thaliana* accessions. This study also delivers insights into how vGWA mapping facilitates the detection and genetic dissection of the genetic architecture of loci contributing to complex traits in natural populations. It thereby illustrates the value of using non-standard statistical methods in genome-wide analyses. Further, it provides an approach to infer allelic heterogeneity which is likely to be both a common, and far too often ignored, complexity in the genetics of multifactorial traits that contributes to undiscovered additive genetic variance and consequently the curse of the “missing heritability”.

## Materials and Methods

### Genotype and Phenotype data

The concentration of molybdenum in leaves was measured in 340 natural *A. thaliana* accessions from the ‘HapMap’ collection ([3]; S1 Table). This dataset contains 58 of the 93 accessions used in the earlier GWA [2] and vGWA [23] analyses of leaf molybdenum concentrations supplemented with 282 newly phenotyped accessions. All accessions were grown in a controlled environment with 6 biological replicate plants per accession, and analyzed by Inductively Coupled Mass Spectroscopy (ICP-MS) for multiple elements including molybdenum, as described previously by Baxter *et al.* [46]. All the ICP-MS data used for the GWA and vGWA is accessible using the digital object identifier (DOI) 10.4231/T9H41PBV, and data for the evaluation of candidate genes using T-DNA insertional alleles is accessible using the DOI 10.4231/ T9SF2T3B (see http://dx.doi.org/).

All accessions have previously been genotyped using the 250k *A. thaliana* SNP chip and that data is publicly available [3]. SNPs where the minor allele frequency was below 5%, were excluded from the analyses. Genotypes were available for more than 95% of the SNPs in all accessions, so none were removed due to problematic genotyping. In total, 200,345 SNPs passed this quality control and were used in our GWA and vGWA analyses.

We evaluated the region upstream of *MOT1* for structural polymorphisms in a set of 283 accessions selected to cover the range of leaf molybdenum concentrations (S4 Table). This was done using gel electrophoresis to identify PCR fragment size differentiation using the primers described in S5 Table. The PCR reactions were completed as follows: 1µl DNA + 5X GoTaq Bf, 2.5mM dNTP’s, 25mM MgCl_2_, 0.4µM of each primer, 0.3µl Taq polymerase, and 9.7µl nuclease free water for a total reaction volume of 25µl. PCR conditions were 94°C for 1 minute to denature, 54°C for 1 minute to anneal, and 72°C for 1.25 minutes for extension, repeated for 40 cycles in the Thermo Px2 thermal cycler (Electron Corporation). DNA was prepared for the accessions that displayed suggestive evidence for structural polymorphisms and submitted for sequencing using Macrogen (dna.macrogen.com). The sequences were then compared to the *Col-0* reference sequence using DiALIGN (http://bibiserv.techfak.uni-bielefeld.de/dialign/) which uncovered five loci and six segregating structural polymorphisms (S2 Table) that were then genotyped in the 283 phenotyped accessions (S4 Table).

### Statistical analyses

All analyses described in the sections below were performed using the R-framework for statistical computing [62].

#### GWA and vGWA analyses

The variance-heterogeneity genome-wide association analyses (vGWA) were performed using the method proposed by Yang *et al.* [22]. In short, this analysis is based on first zscore transforming the analyzed phenotype, and then using the squared zscores for each accession as the phenotype in a normal GWA analysis. The genome scan for genetic effects on the within line variance was performed by using the within-accession coefficient of variation, calculated from the replicate phenotype measurements for each accession, as the phenotype in a standard GWA analysis. The coefficient of variation, rather than the standard deviation, was used as the phenotype since it is a better measure of the within-accession variance due to the removal of the potential scaling effect from differences in the mean phenotype for the accessions. All GWA analyses were performed using a linear mixed model, incorporating the IBS-matrix to correct for population stratification, via the *polygenic* function implemented in the R-package GenABEL [63].

A genome-wide significance threshold was determined for all tested phenotypes by Bonferroni-correction for the number of tested SNPs, resulting in a threshold of 2.5 × 10^-7^. To detect potential inflation of the p-values in the GWA analyses due to remaining population stratification and/or cryptic relatedness, we visually evaluated the relationship between the theoretical distribution of p-values under the null-hypothesis versus those observed in the GWA using quantile-quantile (QQ) plots (S2 Fig), and calculated the inflation factor using the function *estlambda* in the GenABEL package.

#### Multi-locus LASSO regression analyses

Multi-locus regression analysis to identify independent SNP effects on leaf molybdenum concentrations was performed using Lasso regression implemented in the R-package *glmnet* [57]. The Lasso analysis identifies the linear model that minimizes the following 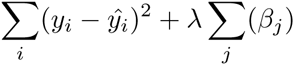 where y_i_ and 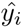 is the phenotype and the predicted phenotype of individual *i*. *β*_*j*_ is the individual genotype effects. The constraint will force most genotype effects to zero, thereby identifying a small subset of polymorphisms with strong independent effects on the phenotype. As λ decreases, the number of non-zero estimates will increase. If λ is zero, the method is identical to an ordinary linear regression. Here, we empirically selected a λ where all SNPs with non-zero effects reached the genome-wide significance threshold in the GWA or vGWA analysis (S3 Fig).

#### DGLM analyses to simultaneously estimate mean and variance effects of evidenced loci

The Double Generalized Linear Model (DGLM) framework allows the simultaneous modelling of both dispersion and mean, by fitting separate linear predictors for both [64]. We fitted a DGLM with separate genetic effects for the variance, and for the mean: *Y ∼ N* (*X*_1_ β_1_, *e*^*X*2^ β^2^) where *Y* is the molybdenum levels, *X*_1_ = [*SNP*_1_, *DEL*_1_, *DUP*_1_]^*T*^, *X*_2_ = *SNP*_*i*_ and *i* is the index of the SNP whose variance effect we are estimating. *β*_1_ and *β*_2_ were estimated using maximum likelihood. This allowed us to include evidenced loci as co-factors with mean effect, while redoing the vGWA scan. The model was fitted using the R-package dglm [65] as suggested in [16].

#### Heritability estimates

Every accession in our data was grown with at least 6 replicates plants. The broad sense heritability (H^2^) was calculated using an ANOVA *y* = β_0_ + *accession \** β_*acc*_ + *e*, comparing within and between line variances.

To calculate the narrow sense heritability (h^2^) we fitted a mixed model 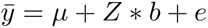 Here 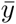 is the mean leaf molybdenum concentration per line and *Z*Z*^*T*^ = *G*, where *G* is the genomic kinship matrix. The intra-class correlation 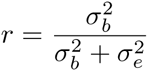 given by this model tells us the amount of variance in 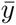 explained by kinship. Assuming that the within line replicates has removed all environmental variance, the amount of the total phenotypic variance explained by kinship, aka h^2^, is *r*H*^2^. In reality, as 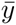 is estimated using < 10 replicates for most lines, some environmental noise will remain in 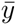, in which case *r * H*^*2*^≤ *h*^*2*^≤ *r.* Here, we therefore present the *r * H*^2^ values, which is the lower bound of h^2^.

#### Variance explained

We estimated the fraction of H^2^ explained by the markers in the *MOT1* region as 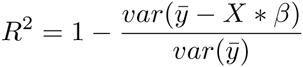, where 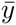 is the mean molybdenum content per line and *X* is the genotype matrix for the markers, fitted as a fixed effect. This estimate assumes that 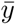 contains no environmental variance which, as stated above, is not entirely the case. If 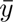 contains environmental noise, this estimate will instead be the lower bound of the fraction of H^2^ explained by *X*, in the same way as described above for h^2^.

The fraction of h^2^ explained by the evaluated set of polymorphisms in the *MOT1* region was estimated by comparing two mixed models:

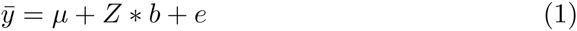

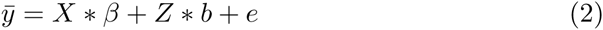

The intra-class correlation *r*_1_ in model (1), gives the amount of variance in 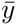 explained by kinship, whereas the intra-class correlation *r*_2_ in model (2) gives the amount of residual variance explained by kinship in this model. To compare the two, we calculate the amount of variance in 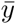 explained by kinship under model (2) as *r*_2,*tot*_ = *r*_2_ * (1 *– R*^*2*^). The fraction of h^2^ explained by the fixed effects *X* are then given as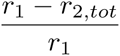. The fraction of variance explained by *X* that is additive is calculated as 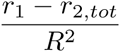.

#### Functional evaluation of candidate genes using T-DNA insertion lines

We identified all genes in the LD-region (r^2^ > 0.4) surrounding the SNP_1_ and SNP_2_ loci. T-DNA insertion lines were ordered for all genes where they were available (Table 3; S3 Table) from the Nottingham Arabidopsis Stock Centre (NASC). Since *MOT1* is known to regulate molybdenum concentrations in *A. thaliana,* the *mot1*-1 T-DNA knockout allele (SALK_118311) for this gene was included on every experimental block of plants as a control, along with *Col-0*. An experimental block is defined by a cultivation tray containing 9 genotypes (including *Col-0* and *mot1-1*) with each genotype represented by between 2 - 12 individuals per block. The average level of replication across all blocks for all genotypes (excluding *Col-0* wild-type and *mot1-1*, which were included in all blocks) was 9.7 individuals (S3 Table). The tested T-DNA insertion lines were grown in 7 independent blocks and the molybdenum concentration in leaves of all plants quantified using the same procedure as used previously [46].

For every experimental block, we compared the molybdenum concentration in leaves between the replicates of every T-DNA insertion line, versus the wild type (*Col-0*), using the non-parametric Wilcox rank test. In 5 out of the 7 blocks, *mot1-1* showed significantly lower molybdenum concentrations compared to the wild type *Col-0* (p < 0.05) as expected, and in one block, the reduction was significant at (p < 0.1). The *mot1-1* mutant in one experimental block of plants showed no difference compared to the wild type, and the results for all other genotypes in this experimental block were therefore discarded.

## Explorations of the long-range LD-block surrounding the *MOT1* gene

The vGWA analyses identify a strong variance-heterogeneity signal across a number of markers in a ∼78 kb segment on chromosome 2 that contains the functional candidate *MOT1* gene (vBLOCK_1_). Visual inspection of the genotype-matrix of this region, sorted by the genotype of the leading SNP in the vGWA analysis (Table 1), indicated the presence of two major groups of accessions that carry the same alleles across a large number of the associated markers (Fig 1C).

## Author contributions

Conceived and designed the study: ÖC DES. Led and coordinated the study: ÖC. Planned and designed the experiments: DES. Planned and designed the computational and quantitative genetic analyses: ÖC SKGF. Contributed reagents/materials/analysis tools: SKGF. Performed the experiments: XH MEA. Analyzed the data: SKGF ÖC XH MEA JD DES. Wrote the paper: SKGF ÖC DES. Commented on the manuscript: MEA, XH, JD.

## Acknowledgements

We thank Lars Rönnegård for advice regarding the statistical analysis, Xia Shen for helpful assistance with providing data, analysis-scripts and input on the data analysis, Mats Pettersson for useful discussions regarding data analysis and interpretations, Yanjun Zan for help preparing figures and useful discussions, and Brett Lahner for the previous ICP-MS analysis.

## Supporting information

**S1 Fig.**
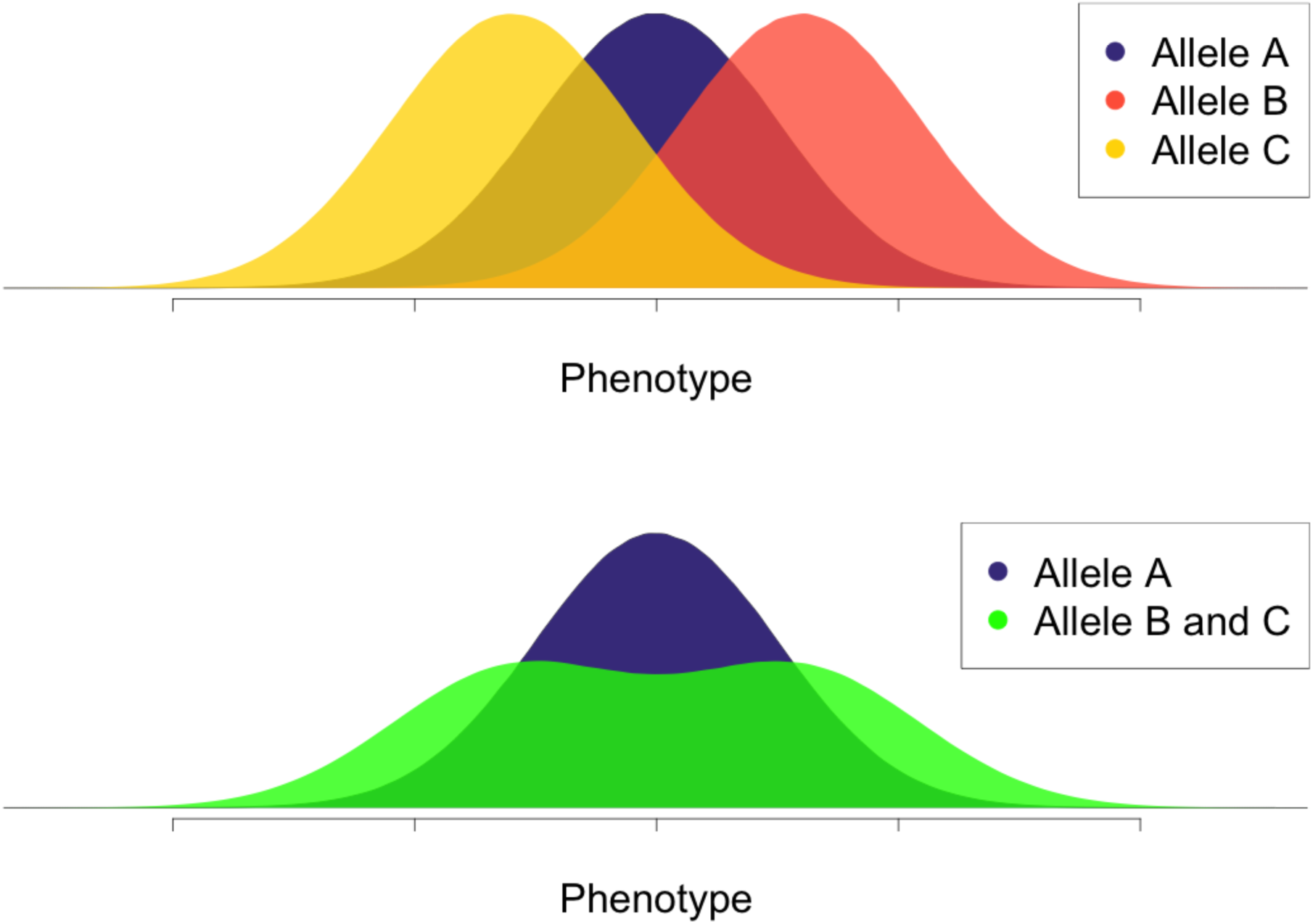
An illustration of how a multi-allelic genetic architecture could lead to variance-heterogeneity signals in a population. The top panel illustrates the hypothetical phenotypic distributions three alleles - A, B and C - that have different effects on a hypothetical trait. The bottom panel illustrate the mixture distributions observed in an association analysis to a bi-allelic marker, where one of the marker-alleles tag functional allele A, and the other tag both alleles B and C. In this situation, no mean difference could be observed between the marker alleles, whereas a large variance difference could be detected via the variance-heterogeneity between the SNP genotypes using a vGWA analysis.

**S2 Fig.**
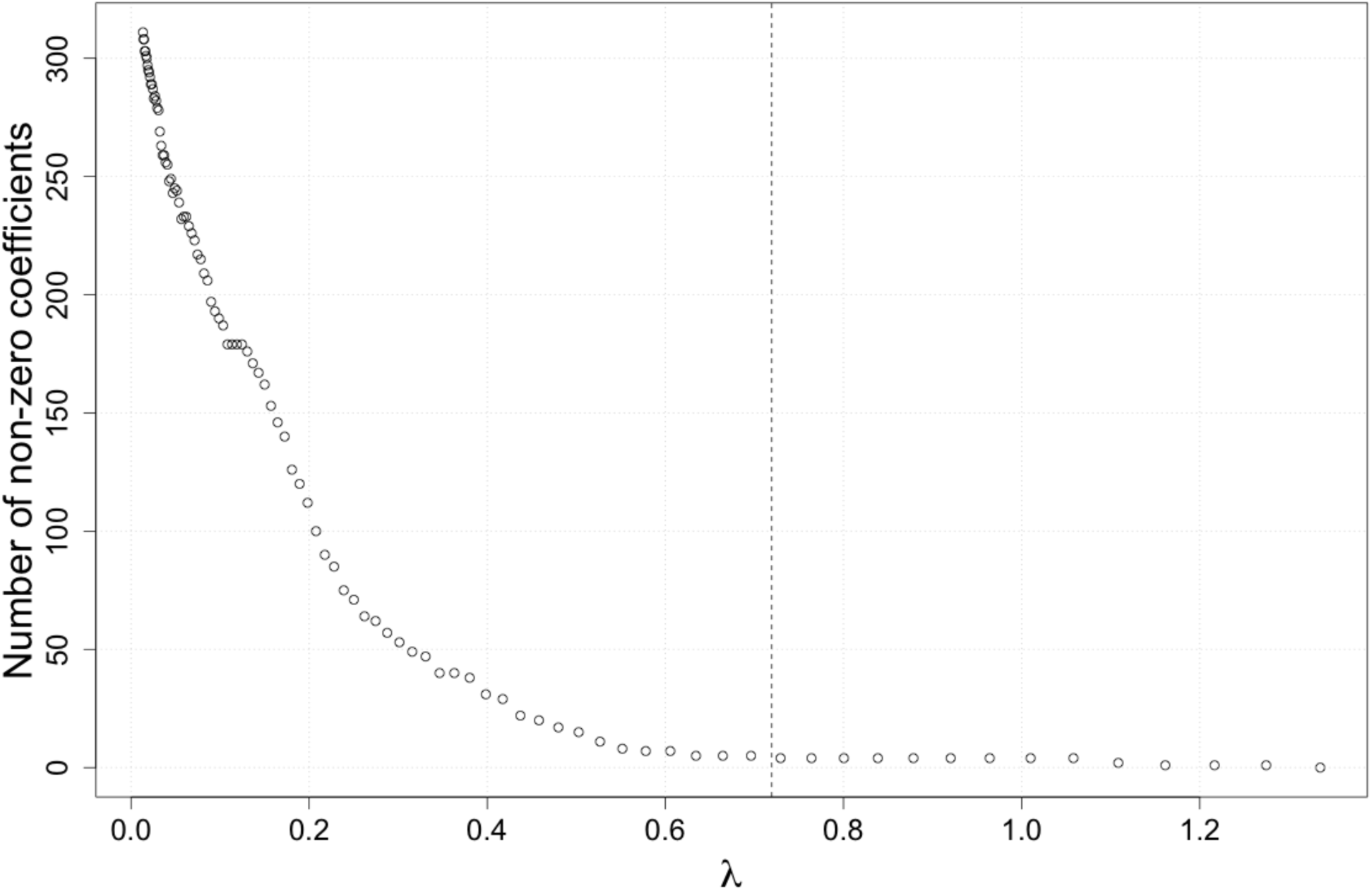
Quantile-quantile (QQ) plots for the genome-wide association analyses to detect genetic effects on the trait mean (GWA), or variance (vGWA). Red line illustrates the theoretical distribution of p-values under the null-hypothesis and the black dots those observed in the two analyses.

**S3 Fig.**
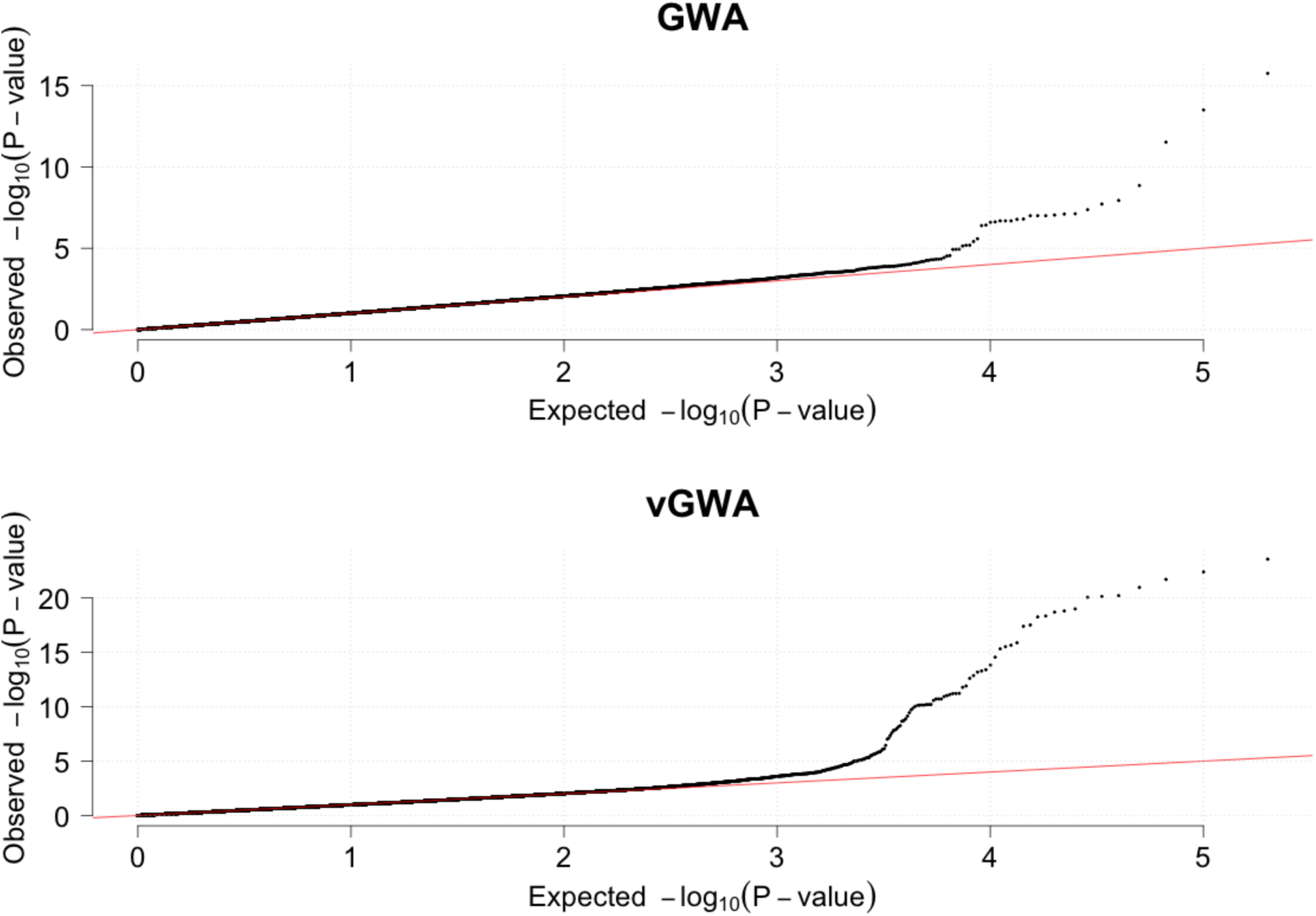
Selection of penalty (λ) in LASSO analysis. Penalty is selected such that all SNPs with non-zero effects in the analysis have reached the genome-wide significance threshold in the GWA or vGWA analysis.

**S1 Table.** Accessions phenotyped for molybdenum content

| **Accession** | **Alternative ID 1** | **Alternative ID 2** | **Replicates** |
| --- | --- | --- | --- |
| LDV.25 | CS76161 | 116 | 6 |
| MOG.37 | CS76189 | 242 | 5 |
| TOU.A1.116 | CS76253 | 282 | 6 |
| LAC.5 | CS76158 | 96 | 6 |
| CAM.16 | CS76107 | 23 | 6 |
| Hi.0 | CS76140 | 8304 | 6 |
| T620 | CS76240 | 6119 | 6 |
| T1110 | CS76236 | 6100 | 6 |
| T1080 | CS76235 | 6098 | 6 |
| TDr.8 | CS76249 | 6194 | 6 |
| Can.0 | CS76109 | 8274 | 6 |
| TOU.E.11 | CS76260 | 366 | 6 |
| UKSE06.482 | CS76289 | 5245 | 6 |
| UKSE06.520 | CS76290 | 5264 | 6 |
| MNF.Che.2 | CS76185 | 1925 | 6 |
| Bur.0 | CS76105 | 5719 | 6 |
| Blh.1 | CS76098 | 7034 | 6 |
| Sparta.1 | CS76229 | 6085 | 6 |
| Hovdala.2 | CS76143 | 6039 | 6 |
| Ull3.4 | CS76295 | 6413 | 6 |
| Rev.2 | CS76219 | 6076 | 4 |
| Fj_1.2 | CS76131 | 6019 | 6 |
| Fj_1.5 | CS76132 | 6020 | 6 |
| Ren.1 | CS76218 | 6959 | 6 |
| Rmx.A180 | CS76220 | 7525 | 6 |
| Se.0 | CS76226 | 6961 | 6 |
| Sq.8 | CS76230 | 6967 | 6 |
| Ull2.3 | CS76293 | 6973 | 6 |
| Ull2.5 | CS76294 | 6974 | 6 |
| Uod.7 | CS76296 | 6976 | 6 |
| NA | CS76298 | 7516 | 6 |
| Wei.0 | CS76301 | 6979 | 6 |
| Ws.0 | CS76303 | 6980 | 6 |
| Wt.5 | CS76304 | 6982 | 6 |
| Yo.0 | CS76305 | 6983 | 6 |
| Zdr.6 | CS76306 | 6985 | 6 |
| Pna.17 | CS76213 | 7523 | 6 |
| Bay.0 | CS76094 | 6899 | 6 |
| Bor.4 | CS76100 | 6903 | 6 |
| Br.0 | CS76101 | 6904 | 6 |
| C24 | CS76106 | 6906 | 6 |
| Col.0 | CS76113 | 6909 | 158 |
| Cvi.0 | CS76116 | 6911 | 159 |
| Est.1 | CS76127 | 6916 | 6 |
| Fei.0 | CS76129 | 8215 | 6 |
| Got.7 | CS76136 | 6921 | 6 |
| Ler.1 | CS76164 | 6932 | 6 |
| L_v.5 | CS76175 | 6046 | 6 |
| NFA.8 | CS76199 | 6944 | 6 |
| RRS.10 | CS22689 | 7515 | 6 |
| RRS.7 | CS28713 | 7514 | 6 |
| Shahdara | CS76227 | 6962 | 6 |
| Tamm.2 | CS76244 | 6968 | 6 |
| Ts.1 | CS76268 | 6970 | 159 |
| Van.0 | CS76297 | 6977 | 6 |
| Lm.2 | CS76173 | 8329 | 6 |
| Uk.2 | CS28788 | 7379 | 6 |
| Co.4 | CS28165 | 7080 | 6 |
| Lab.5 | CS28181 | 6744 | 6 |
| Di.1 | CS28208 | 7098 | 6 |
| Gr.5 | CS28326 | 7158 | 6 |
| Ha.0 | CS28336 | 7163 | 6 |
| Hn.0 | CS28350 | 7165 | 6 |
| Sav.0 | CS28725 | 7340 | 6 |
| Tscha.1 | CS28779 | 7372 | 6 |
| Wa.1 | CS28804 | 7394 | 6 |
| Ws | CS28823 | 7397 | 6 |
| TOU.A1.115 | CS76252 | 281 | 6 |
| Fi.1 | CS28252 | 7139 | 6 |
| Pn.0 | CS28645 | 7307 | 6 |
| Sh.0 | CS28734 | 7331 | 6 |
| Si.0 | CS28739 | 7337 | 5 |
| Uk.1 | CS28787 | 7378 | 6 |
| Wc.2 | CS28814 | 7405 | 6 |
| CAM.61 | CS76108 | 66 | 6 |
| LDV.58 | CS76163 | 149 | 6 |
| TOU.A1.62 | CS76256 | 328 | 6 |
| SLSP.30 | CS76228 | 2274 | 6 |
| UKID48 | CS76273 | 5753 | 6 |
| Bg.2 | CS76096 | 6709 | 6 |
| Aa.0 | CS28007 | 7000 | 6 |
| Alst.1 | CS28013 | 6989 | 6 |
| Bsch.0 | CS28099 | 7031 | 6 |
| Ca.0 | CS28128 | 7062 | 4 |
| Da(1).12 | CS28201 | 7460 | 6 |
| Ep.0 | CS28236 | 7123 | 6 |
| Est.0 | CS28243 | 7128 | 6 |
| Ge.1 | CS28277 | 7145 | 6 |
| Gie.0 | CS28280 | 7147 | 6 |
| Hey.1 | CS28344 | 7166 | 6 |
| Mh.0 | CS28492 | 7255 | 6 |
| No.0 | CS28564 | 7275 | 6 |
| Nw.0 | CS28573 | 7258 | 6 |
| Pa.2 | CS28595 | 7291 | 6 |
| Pr.0 | CS28651 | 7310 | 6 |
| Sapporo.0 | CS28724 | 7330 | 6 |
| Sei.0 | CS28729 | 7333 | 6 |
| Wl.0 | CS28822 | 7411 | 6 |
| MIB.28 | CS76183 | 178 | 6 |
| TOU.I.17 | CS76263 | 378 | 6 |
| LI.OF.095 | CS76166 | 8241 | 6 |
| Alc.0 | CS76088 | 6988 | 6 |
| NA | CS76093 | 8256 | 6 |
| 11PNA4.101 | CS76084 | 8796 | 6 |
| Br_1.6 | CS76102 | 8231 | 6 |
| Bu.0 | CS76103 | 8271 | 6 |
| TDr.3 | CS76248 | 6190 | 6 |
| Cen.0 | CS76110 | 8275 | 6 |
| Kelsterbach.4 | CS76152 | 8420 | 6 |
| Dra3.1 | CS76117 | 8283 | 6 |
| Drall.1 | CS76118 | 8284 | 6 |
| Duk | CS76124 | 6008 | 6 |
| Fj_1.1 | CS76130 | 8422 | 6 |
| Gd.1 | CS76134 | 8296 | 6 |
| Ge.0 | CS76135 | 8297 | 6 |
| Gr.1 | CS76137 | 8300 | 6 |
| Hod | CS76141 | 8235 | 6 |
| Hov4.1 | CS76142 | 8306 | 6 |
| Hs.0 | CS76145 | 8310 | 6 |
| HSm | CS76146 | 8236 | 6 |
| In.0 | CS76147 | 8311 | 6 |
| Ka.0 | CS76149 | 8314 | 6 |
| K_In | CS76155 | 8239 | 6 |
| Kulturen.1 | CS76156 | 8240 | 6 |
| Lc.0 | CS76159 | 8323 | 6 |
| Liarum | CS76167 | 8242 | 6 |
| Lip.0 | CS76168 | 8325 | 6 |
| Lis.1 | CS76169 | 8326 | 6 |
| Lis.2 | CS76170 | 8222 | 6 |
| Lisse | CS76171 | 8430 | 6 |
| Lom1.1 | CS76174 | 6042 | 6 |
| Lund | CS76178 | 8335 | 6 |
| Na.1 | CS76195 | 8343 | 6 |
| NA | CS76201 | 6074 | 6 |
| Ost.0 | CS76202 | 8351 | 6 |
| Pa.1 | CS76204 | 8353 | 6 |
| Per.1 | CS76210 | 8354 | 6 |
| Petergof | CS76211 | 7296 | 6 |
| Rak.2 | CS76217 | 8365 | 6 |
| Rsch.4 | CS76222 | 8374 | 6 |
| Sanna.2 | CS76223 | 8376 | 6 |
| Sap.0 | CS76224 | 8378 | 6 |
| Sav.0 | CS76225 | 8412 | 6 |
| St.0 | CS76231 | 8387 | 6 |
| Ta.0 | CS76242 | 8389 | 6 |
| Tottarp.2 | CS76251 | 6243 | 6 |
| PHW.36 | CS28636 | 7507 | 6 |
| Sp.0 | CS28743 | 7343 | 6 |
| DraIV.6.35 | CS76123 | 6005 | 6 |
| UKID22 | CS76271 | 5729 | 6 |
| UKNW06.059 | CS76275 | 5380 | 6 |
| UKNW06.060 | CS76276 | 5381 | 6 |
| UKNW06.386 | CS76277 | 5565 | 6 |
| Be.1 | CS28063 | 7011 | 6 |
| Kl.5 | CS28394 | 7199 | 6 |
| Li.3 | CS28454 | 7224 | 6 |
| Ob.1 | CS28580 | 7277 | 6 |
| PHW.20 | CS28620 | 7490 | 6 |
| PHW.22 | CS28622 | 7492 | 6 |
| Pla.0 | CS28640 | 7300 | 6 |
| Pog.0 | CS28650 | 7306 | 6 |
| Wt.3 | CS28833 | 7408 | 6 |
| Zu.1 | CS28847 | 7418 | 6 |
| DraIV.1.5 | CS76120 | 5887 | 6 |
| DraIV.6.16 | CS76122 | 5987 | 6 |
| MIB.22 | CS76182 | 173 | 6 |
| TOU.A1.96 | CS76258 | 357 | 6 |
| PAR.3 | CS76205 | 258 | 6 |
| TOU.H.13 | CS76262 | 374 | 6 |
| LAC.3 | CS76157 | 94 | 6 |
| MNF.Pot.48 | CS76187 | 1859 | 6 |
| TDr.1 | CS76245 | 6188 | 6 |
| UKSE06.429 | CS76287 | 5207 | 2 |
| Zdrl.2.24 | CS76307 | 6448 | 6 |
| Ba.1 | CS28053 | 7014 | 6 |
| Blh.2 | CS28090 | 7035 | 6 |
| Knox.11 | CS28407 | 6810 | 6 |
| PHW.28 | CS28628 | 7498 | 6 |
| PHW.35 | CS28635 | 7506 | 6 |
| Wag.3 | CS28808 | 7390 | 6 |
| Ors.2 | CS28849 | 7284 | 6 |
| Paw.3 | CS76208 | 2150 | 6 |
| Wil.1 | CS76302 | 100000 | 6 |
| Ag.0 | CS76087 | 6897 | 6 |
| An.1 | CS76091 | 6898 | 6 |
| Bor.1 | CS76099 | 5837 | 6 |
| CIBC.17 | CS76111 | 6907 | 6 |
| N13 | CS76194 | 7438 | 6 |
| Ct.1 | CS76114 | 6910 | 6 |
| Eden.2 | CS76125 | 6913 | 6 |
| Edi.0 | CS76126 | 6914 | 6 |
| Ga.0 | CS76133 | 6919 | 6 |
| Gy.0 | CS76139 | 8214 | 6 |
| HR.5 | CS76144 | 6924 | 6 |
| Kas.2 | CS76150 | 8424 | 6 |
| Kin.0 | CS76153 | 6926 | 6 |
| Kno.18 | CS76154 | 6928 | 6 |
| LL.0 | CS76172 | 6933 | 6 |
| Lp2.2 | CS76176 | 7520 | 6 |
| Lp2.6 | CS76177 | 7521 | 6 |
| Lz.0 | CS76179 | 6936 | 6 |
| Mr.0 | CS76190 | 7522 | 6 |
| Mrk.0 | CS76191 | 6937 | 6 |
| Mt.0 | CS76192 | 6939 | 6 |
| Mz.0 | CS76193 | 6940 | 6 |
| Nd.1 | CS76197 | 6942 | 6 |
| NFA.10 | CS76198 | 6943 | 6 |
| Oy.0 | CS76203 | 6946 | 6 |
| Pro.0 | CS76214 | 8213 | 6 |
| Pu2.23 | CS76215 | 6951 | 6 |
| Ra.0 | CS76216 | 6958 | 6 |
| Bla.1 | CS76097 | 7015 | 5 |
| Cha.0 | CS28133 | 7069 | 6 |
| PHW.31 | CS28631 | 7502 | 6 |
| Sei.0 | CS28732 | 7344 | 6 |
| Ste.0 | CS28750 | 7346 | 6 |
| Tsu.0 | CS28780 | 7373 | 6 |
| Ven.1 | CS28800 | 7384 | 6 |
| CUR.3 | CS76115 | 81 | 6 |
| TOU.H.12 | CS76261 | 373 | 6 |
| TOU.J.3 | CS76266 | 383 | 6 |
| MNF.Pot.68 | CS76188 | 1867 | 6 |
| Belmonte.4.94 | CS76095 | 957 | 6 |
| An.2 | CS28017 | 6996 | 6 |
| Benk.1 | CS28064 | 7008 | 6 |
| CIBC4 | CS28141 | 6729 | 6 |
| Com.1 | CS28193 | 7092 | 6 |
| Hh.0 | CS28345 | 7169 | 6 |
| Je.0 | CS28364 | 7181 | 6 |
| Mc.0 | CS28490 | 7252 | 6 |
| N4 | CS28510 | 7446 | 6 |
| Nok.1 | CS28568 | 7270 | 6 |
| PHW.14 | CS28614 | 7483 | 6 |
| Rhen.1 | CS28685 | 7316 | 6 |
| Ty.0 | CS28786 | 7351 | 6 |
| Wag.4 | CS28809 | 7391 | 6 |
| ALL1.2 | CS76089 | 1 | 6 |
| VOU.2 | CS76300 | 392 | 6 |
| TOU.I.2 | CS76264 | 379 | 6 |
| TOU.I.6 | CS76265 | 380 | 6 |
| ROM.1 | CS76221 | 267 | 6 |
| Map.42 | CS76180 | 2057 | 6 |
| UKID37 | CS76272 | 5742 | 5 |
| UKSE06.192 | CS76281 | 5056 | 6 |
| UKSE06.278 | CS76283 | 5122 | 6 |
| UKSE06.349 | CS76284 | 5158 | 6 |
| App1.16 | CS76092 | 5832 | 6 |
| Ann.1 | CS28049 | 6994 | 6 |
| Baa.1 | CS28054 | 7002 | 6 |
| Boot.1 | CS28091 | 7026 | 6 |
| CIBC5 | CS28142 | 6730 | 6 |
| Cit.0 | CS28158 | 7075 | 6 |
| Es.0 | CS28241 | 7126 | 6 |
| Li.5:2 | CS28457 | 7227 | 6 |
| Li.6 | CS28459 | 7229 | 6 |
| N7 | CS28513 | 7449 | 6 |
| NFC20 | CS28550 | 6847 | 6 |
| PUZ24 | CS28663 | 6953 | 6 |
| Rou.0 | CS28692 | 7320 | 6 |
| Ting.1 | CS28759 | 7354 | 6 |
| ALL1.3 | CS76090 | 2 | 6 |
| TOU.K.3 | CS76267 | 386 | 6 |
| KBS.Mac.8 | CS76151 | 1716 | 6 |
| MNF.Jac.32 | CS76186 | 1967 | 6 |
| UKID80 | CS76274 | 5785 | 6 |
| UKSW06.202 | CS76292 | 4802 | 6 |
| UKSE06.272 | CS76282 | 5116 | 5 |
| UKSE06.414 | CS76286 | 5202 | 6 |
| Chat.1 | CS28135 | 7071 | 6 |
| Co.2 | CS28163 | 7078 | 6 |
| Da.0 | CS28200 | 7094 | 6 |
| Ga.2 | CS28274 | 7141 | 6 |
| Gel.1 | CS28279 | 7143 | 6 |
| Gì\|.0 | CS28282 | 7151 | 6 |
| NA | CS28332 | 7150 | 6 |
| Jl.3 | CS28369 | 7424 | 6 |
| Jm.1 | CS28373 | 7178 | 6 |
| Kelsterbach.2 | CS28382 | 7188 | 6 |
| Kr.0 | CS28419 | 7201 | 6 |
| Kro.0 | CS28420 | 7206 | 6 |
| Krot.2 | CS28423 | 7205 | 6 |
| Li.7 | CS28461 | 7231 | 6 |
| Mnz.0 | CS28495 | 7244 | 6 |
| Nw.2 | CS28575 | 7260 | 6 |
| Nz1 | CS28578 | 7263 | 6 |
| Old.1 | CS28583 | 7280 | 6 |
| Or.0 | CS28587 | 7282 | 6 |
| S96 | CS28720 | 7472 | 6 |
| Utrecht | CS28795 | 7382 | 6 |
| Wag.5 | CS28810 | 7392 | 6 |
| WAR | CS28812 | 7477 | 6 |
| 11ME1.32 | CS76083 | 8610 | 6 |
| 328PNA054 | CS76085 | 8692 | 6 |
| CIBC2 | CS28140 | 6727 | 6 |
| Dra.2 | CS28214 | 7105 | 6 |
| PHW.10 | CS28610 | 7479 | 6 |
| PHW.13 | CS28613 | 7482 | 6 |
| PHW.33 | CS28633 | 7504 | 6 |
| PHW.37 | CS28637 | 7508 | 6 |
| Tiv.1 | CS28760 | 7355 | 6 |
| DraIV.1.14 | CS76119 | 5896 | 6 |
| MIB.15 | CS76181 | 166 | 6 |
| MIB.84 | CS76184 | 223 | 6 |
| LDV.34 | CS76162 | 126 | 6 |
| VOU.1 | CS76299 | 390 | 6 |
| TOU.A1.43 | CS76255 | 321 | 6 |
| PAR.4 | CS76206 | 259 | 6 |
| TOU.C.3 | CS76259 | 362 | 6 |
| PAR.5 | CS76207 | 260 | 6 |
| JEA | CS76148 | 91 | 6 |
| Lill_.1 | CS76165 | 641 | 6 |
| UKSE06.351 | CS76285 | 5160 | 6 |
| UKSE06.466 | CS76288 | 5232 | 6 |
| UKNW06.436 | CS76278 | 5606 | 6 |
| UKNW06.460 | CS76279 | 5628 | 6 |
| Bs.2 | CS28097 | 7004 | 6 |
| Db.0 | CS28202 | 7100 | 6 |
| Do.0 | CS28210 | 7102 | 6 |
| Ede.1 | CS28217 | 7110 | 6 |
| Fr.4 | CS28268 | 7135 | 6 |
| Kn.0 | CS28395 | 7186 | 6 |
| Nc.1 | CS28527 | 7430 | 5 |
| Pent.1 | CS76209 | 2187 | 6 |
| T1040 | CS76233 | 6094 | 6 |
| T1060 | CS76234 | 6096 | 6 |
| T510 | CS76238 | 6109 | 6 |
| T540 | CS76239 | 6112 | 6 |
| T690 | CS76241 | 6124 | 5 |
| NA | CS76243 | 6169 | 6 |
| TDr.17 | CS76246 | 6202 | 6 |
| TDr.18 | CS76247 | 6203 | 6 |
| Tomegap.2 | CS76250 | 6242 | 6 |
| Udul.1.34 | CS76269 | 6318 | 6 |
| Amel.1 | CS28014 | 6990 | 6 |
| Ang.0 | CS28018 | 6992 | 6 |
| Ste.3 | CS76232 | 2290 | 6 |
| UKID101 | CS76270 | 5805 | 6 |
| UKSE06.062 | CS76280 | 4997 | 6 |
| UKSE06.628 | CS76291 | 5341 | 5 |
| Zdrl.2.25 | CS76308 | 6449 | 6 |

**S2 Table.** Structural polymorphisms in *MOT1* promoter region

| Type | Locus | Allele<sup>1</sup> | Length (bp) | Position<sup>2</sup> |
| --- | --- | --- | --- | --- |
| Deletion | DEL<sub>1</sub> | DEL<sub>1</sub><sup>53</sup> | 53 | 10934564 |
| Deletion | DEL<sub>2</sub> | DEL<sub>2</sub><sup>49</sup> | 49 | 10935270 |
| Insertion | INS<sub>1</sub> | INS<sub>1</sub><sup>14</sup> | 14 | 10936794 |
| Insertion | INS<sub>2</sub> | INS<sub>2</sub><sup>39</sup> | 39 | 10936780 |
| Duplication | DUP<sub>1</sub> | DUP<sub>1</sub><sup>326</sup> | 326 | 10934983 |
| Duplication | DUP<sub>1</sub> | DUP<sub>1</sub><sup>322</sup> | 322 | 10934945 |

<sup>1</sup>Abbreviation for minor (structural) allele
<sup>2</sup>Location (bp) on Chromosome 2, TAIR10 assembly

**S3 Table.**
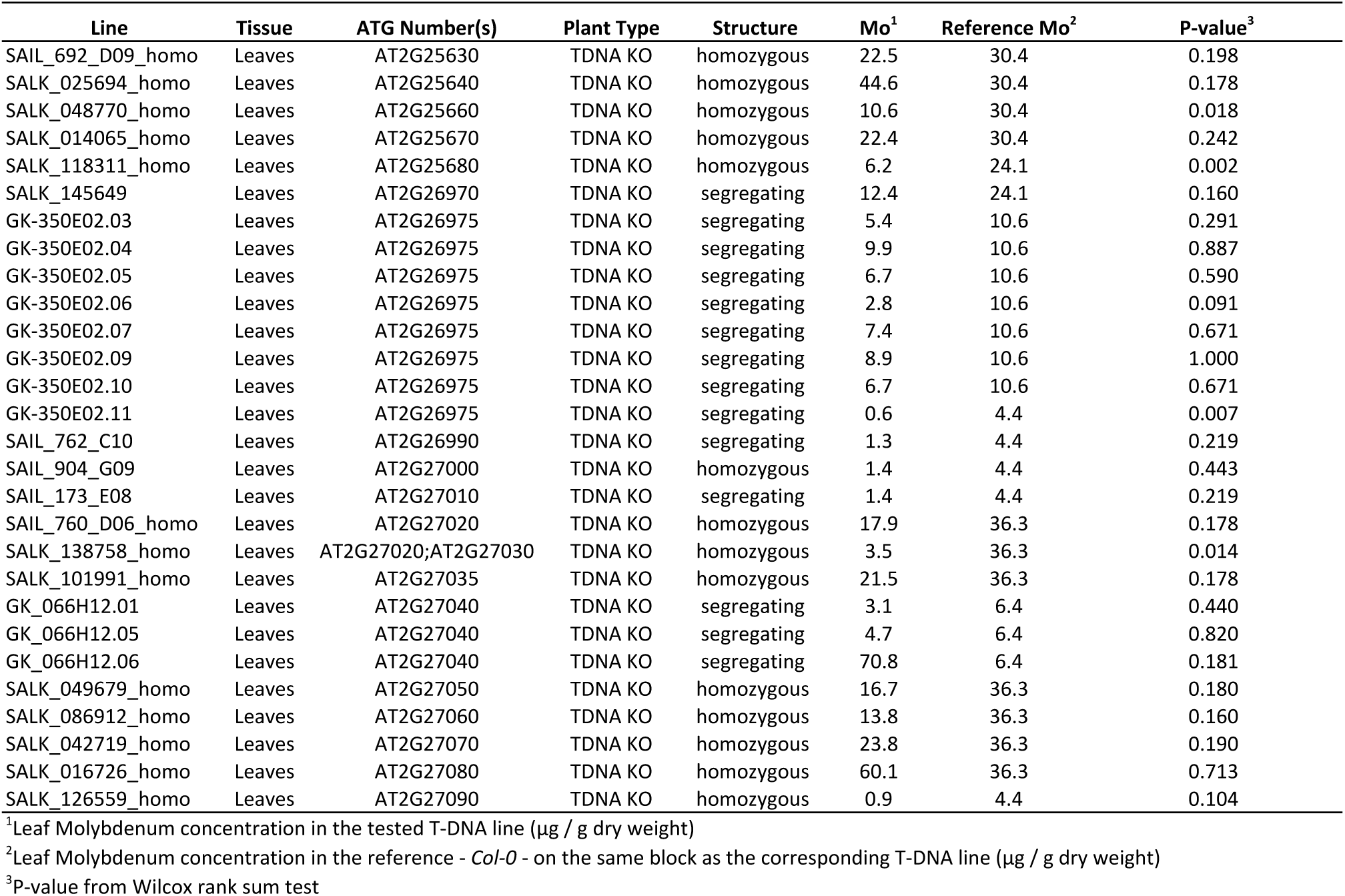
Tested T-DNA insertion lines

**S4 Table.**
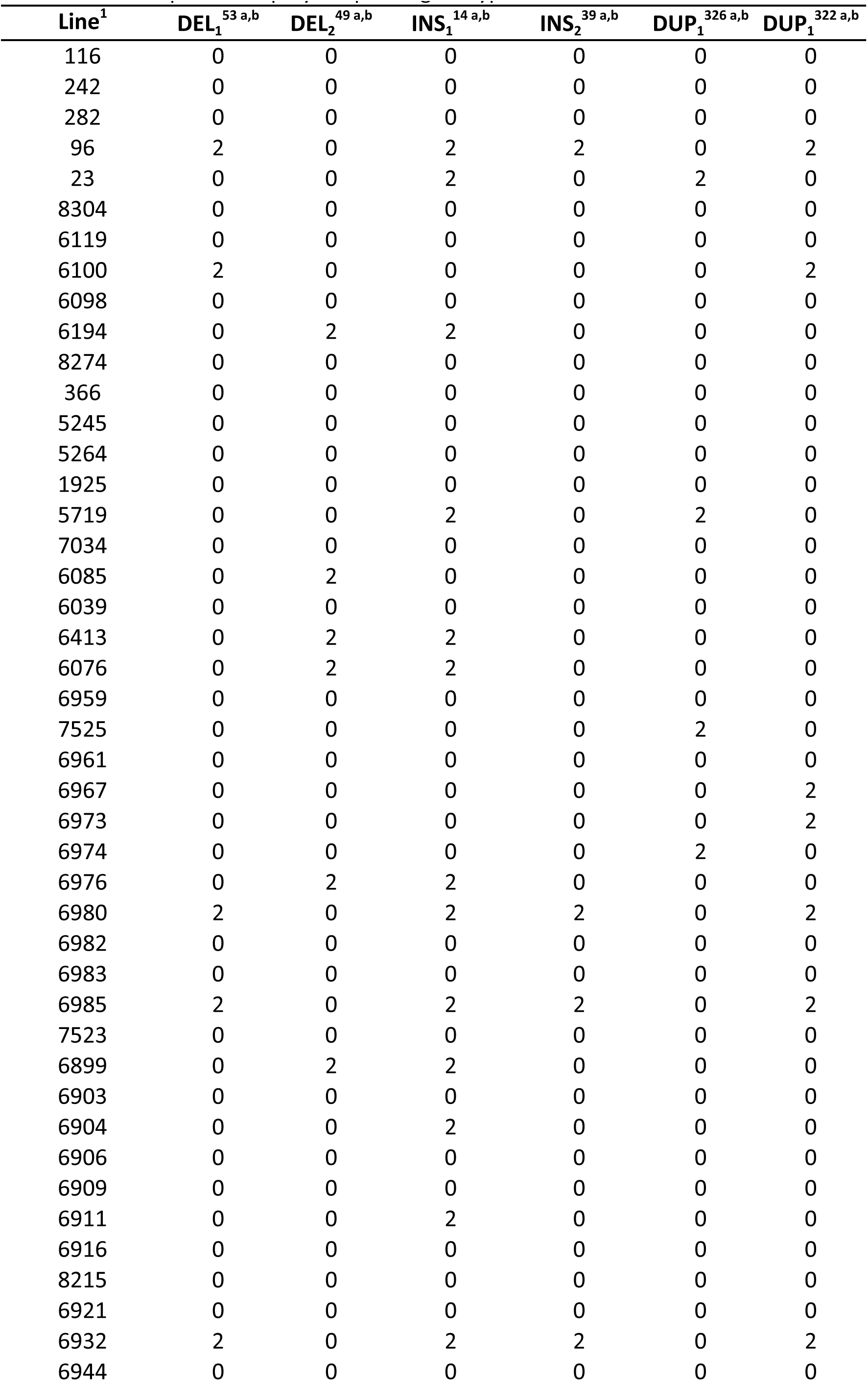
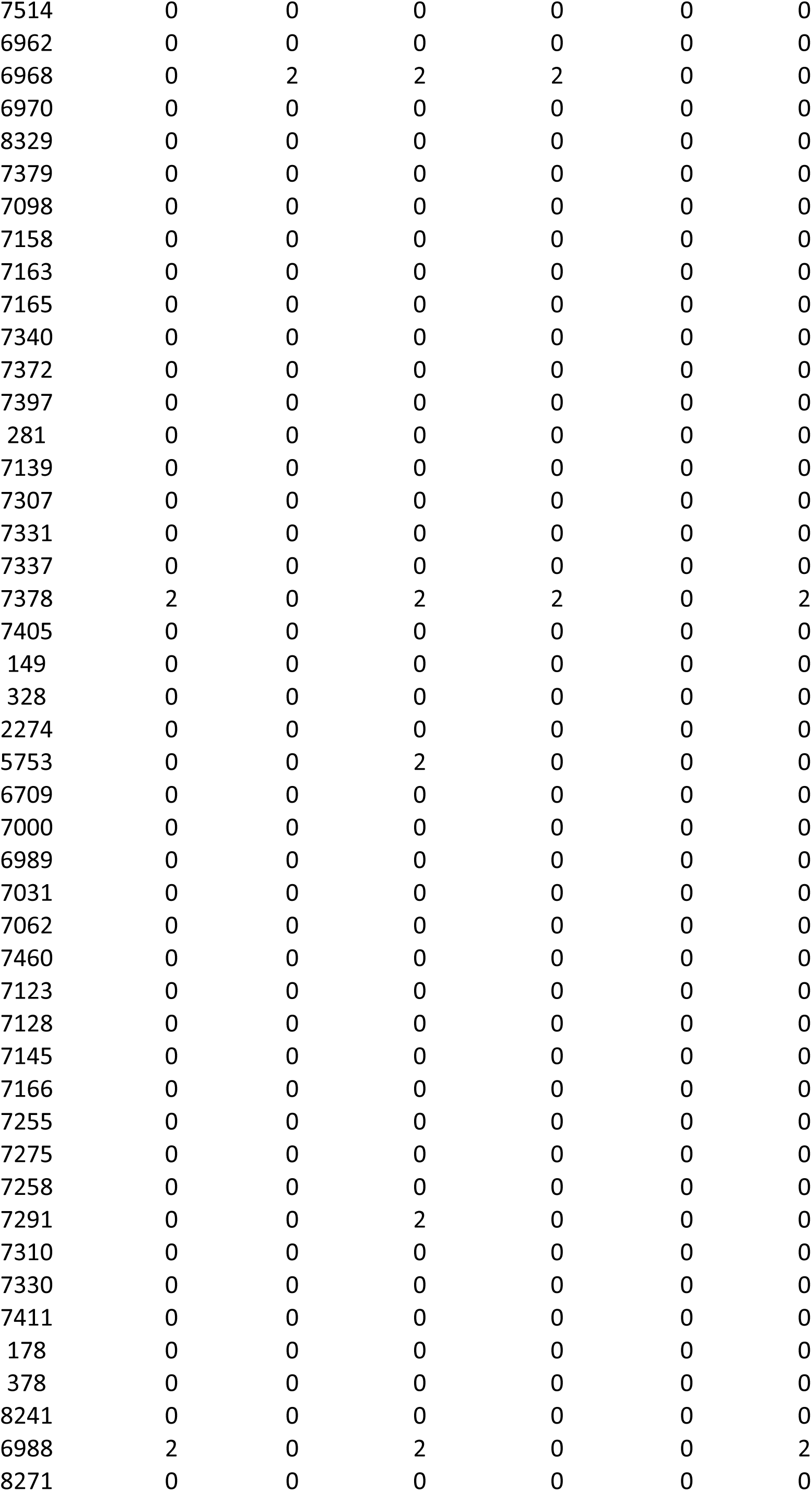
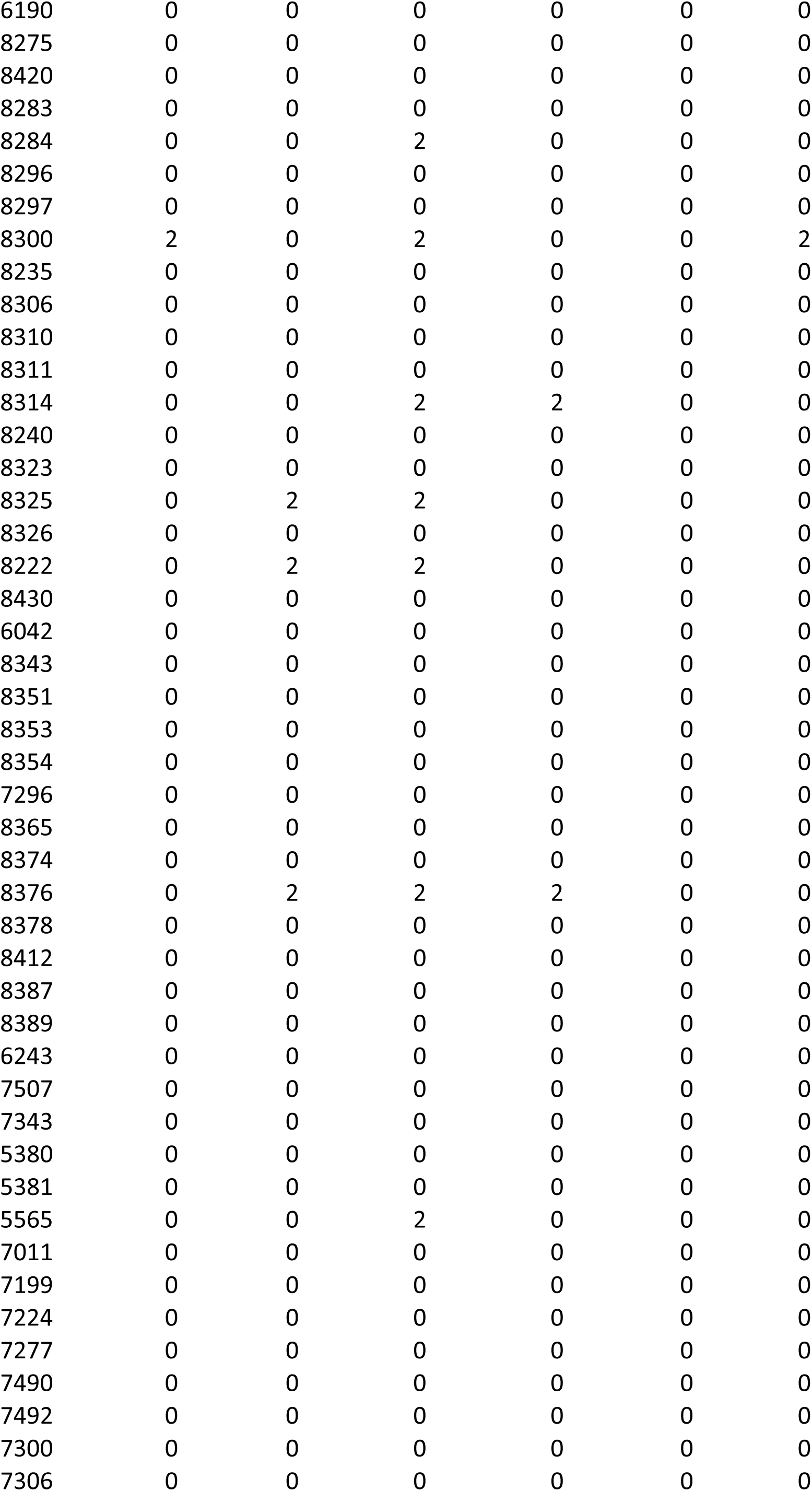
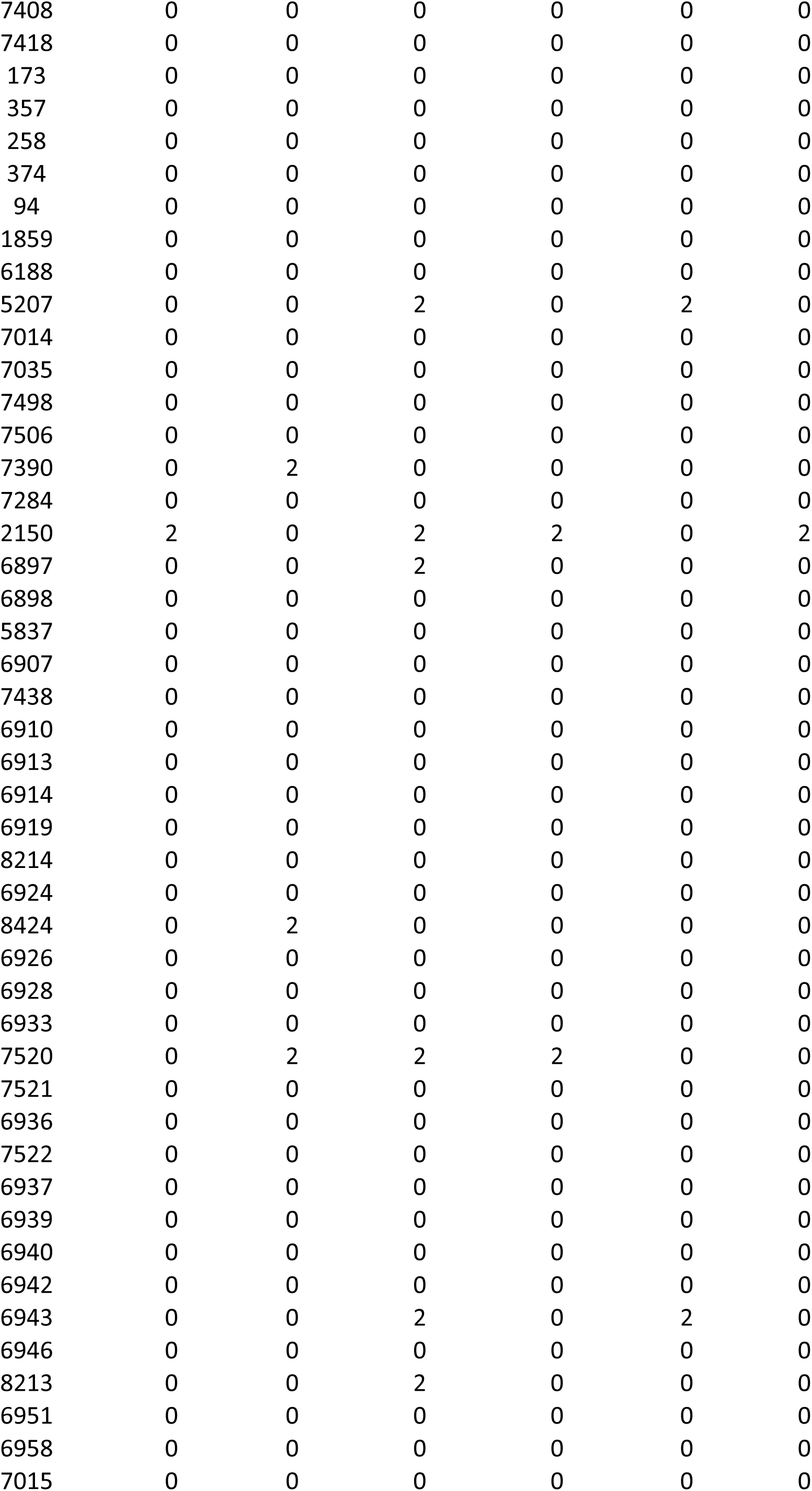
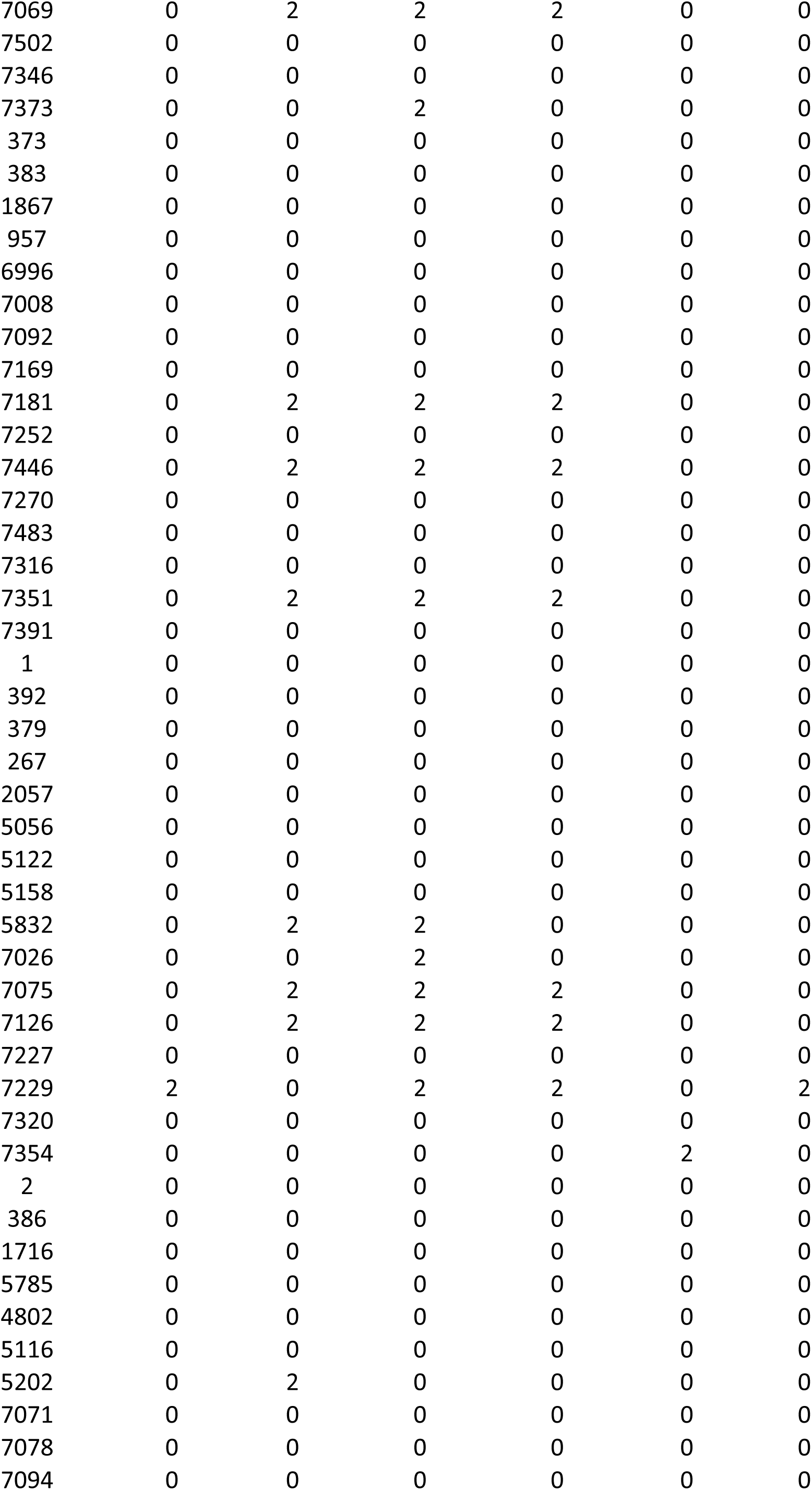
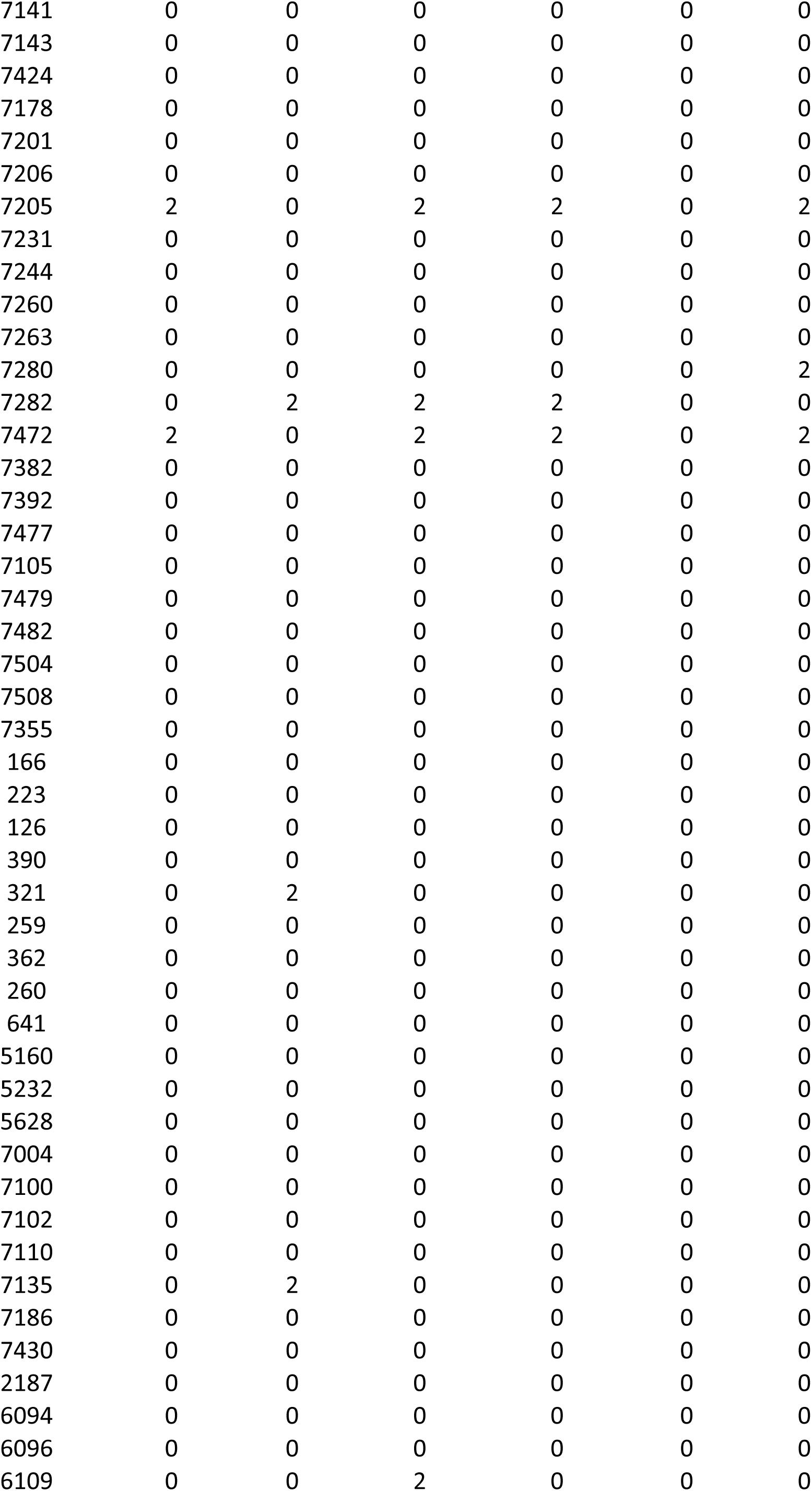
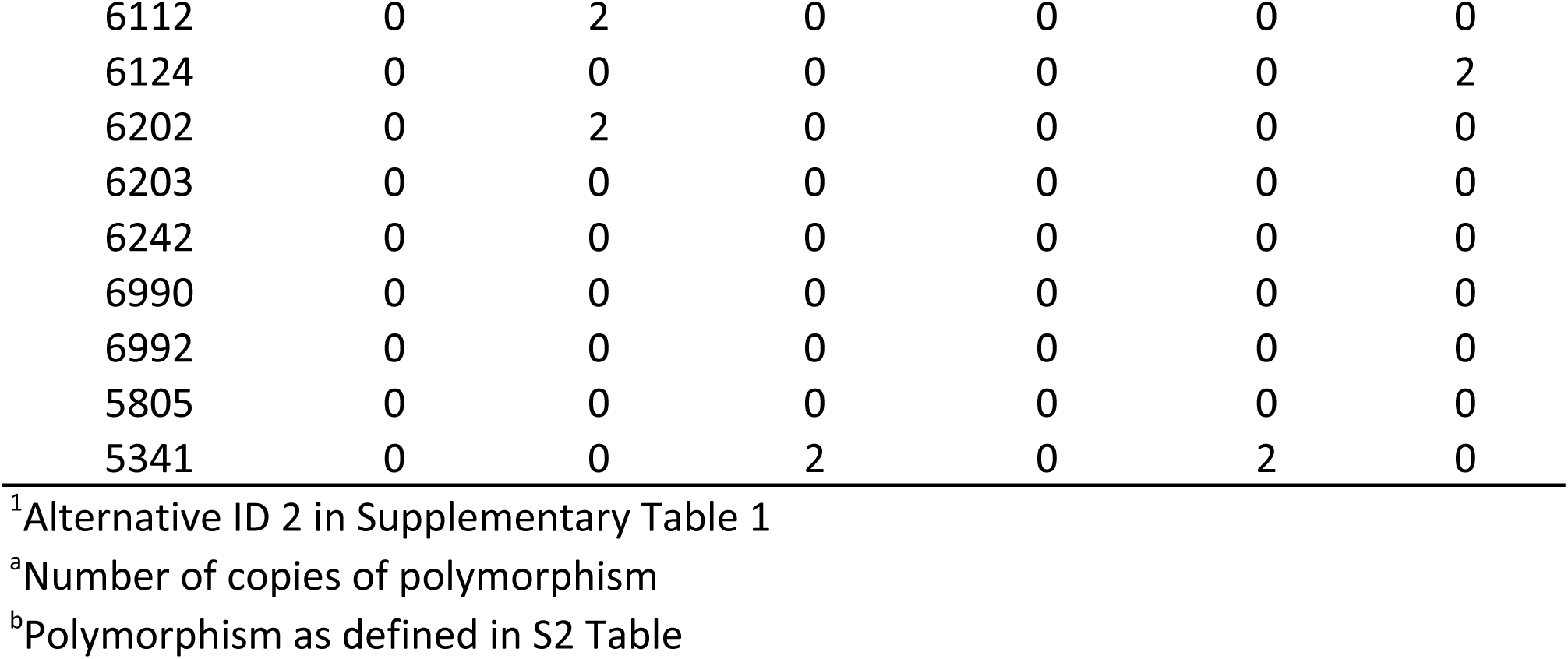
*MOT1* promoter polymorphism genotypes for 283 *A. thaliana* accessions.

**S5 Table.**
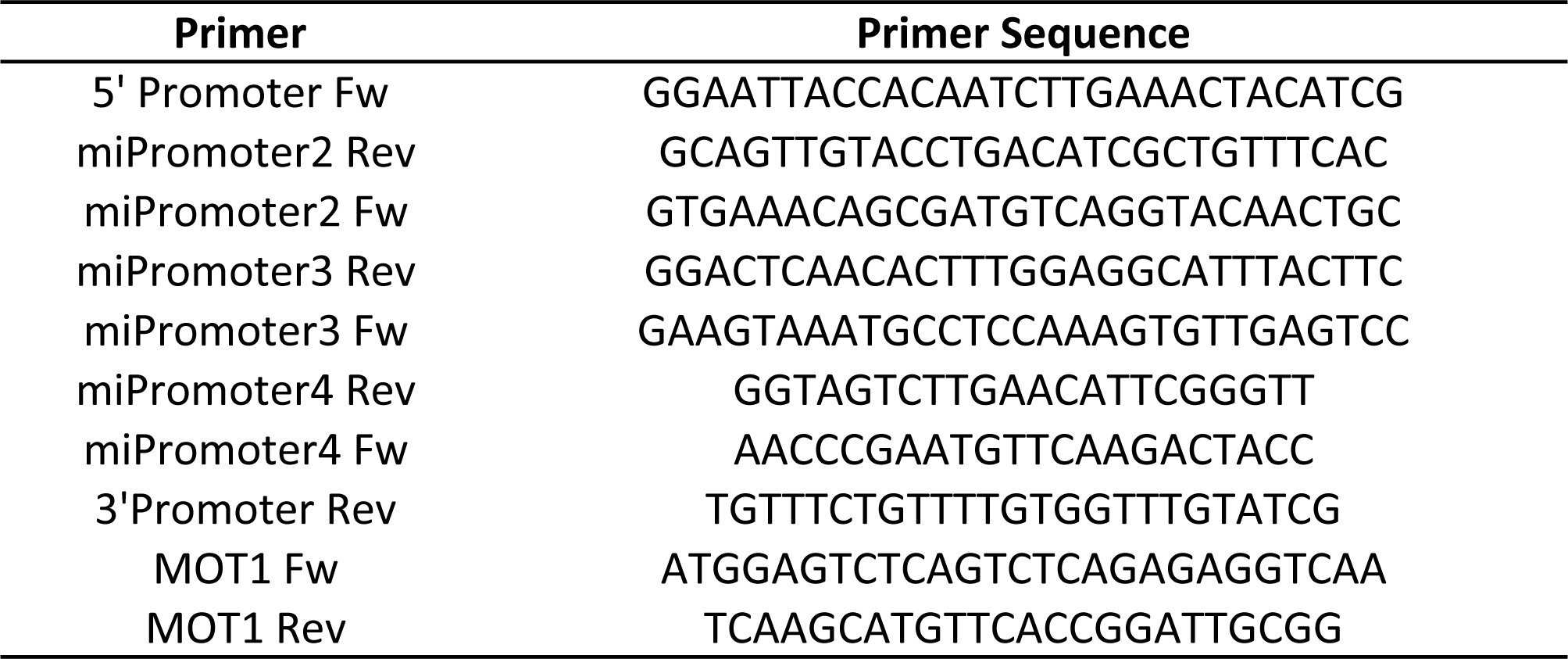
Primers for genotyping the promotor region of *MOT1*.

